# A comprehensive analysis of gene expression changes in a high replicate and open-source dataset of differentiating hiPSC-derived cardiomyocytes

**DOI:** 10.1101/2021.04.22.441027

**Authors:** Tanya Grancharova, Kaytlyn A. Gerbin, Alexander B. Rosenberg, Charles M. Roco, Joy Arakaki, Colette DeLizzo, Stephanie Q. Dinh, Rory Donovan-Maiye, Matthew Hirano, Angelique Nelson, Joyce Tang, Julie A. Theriot, Calysta Yan, Vilas Menon, Sean P. Palecek, Georg Seelig, Ruwanthi N. Gunawardane

## Abstract

We performed a comprehensive analysis of the transcriptional changes within and across cell populations during human induced pluripotent stem cell (hiPSC) differentiation to cardiomyocytes. Using the single cell RNA-seq combinatorial barcoding method SPLiT-seq, we sequenced >20,000 single cells from 55 independent samples representing two differentiation protocols and multiple hiPSC lines. Samples included experimental replicates ranging from undifferentiated hiPSCs to mixed populations of cells at D90 post-differentiation. As expected, differentiated cell populations clustered by time point, with differential expression analysis revealing markers of cardiomyocyte differentiation and maturation changing from D12 to D90. We next performed a complementary cluster-independent sparse regression analysis to identify and rank genes that best assigned cells to differentiation time points. The two highest ranked genes between D12 and D24 (*MYH7* and *MYH6*) resulted in an accuracy of 0.84, and the three highest ranked genes between D24 and D90 (*A2M, H19, IGF2*) resulted in an accuracy of 0.94, revealing that low dimensional gene features can identify differentiation or maturation stages in differentiating cardiomyocytes. Expression levels of select genes were validated using RNA FISH. Finally, we interrogated differences in differentiation population composition and cardiac gene expression resulting from two differentiation protocols, experimental replicates, and three hiPSC lines in the WTC-11 background to identify sources of variation across these experimental variables.

## Introduction

Single cell RNA-sequencing (scRNA-seq) has revolutionized the study of heterogeneous cell populations and cell state transitions at the single cell level, including during differentiation and development. ScRNA-seq datasets enable identification and characterization of transcriptionally distinct cell types and transitions *in vivo* and *in vitro* by revealing the expression of cell type-specific genes or gene modules^1-5^. This has allowed for the identification of unique cell types and transient states, including rare types that were previously undetectable by bulk sequencing methods^4,6-8^.

Cell state transitions during the differentiation of human pluripotent stem cells to cardiomyocytes have also been profiled by scRNA-seq^9,10^ and are characterized by some of the same gene expression programs found during *in vivo* development^6,11-13^. Transcriptional profiling of *in vitro*-derived cardiomyocytes after extended time in culture or after perturbations can serve as a benchmark, classifying maturation states and marker genes that are associated with cell function and phenotype^14-16^. Identifying cardiac maturation state is important in settings where the utility of *in vitro* derived cardiomyocytes is limited by their immaturity and heterogeneity, such as *in vivo* cell transplantation where immature cells can be pro-arrhythmogenic^17-19^, and drug testing and phenotyping platforms where the degree of functional maturation affects cell performance^14,20^. Despite the importance of characterizing populations of more mature cardiomyocytes, most scRNA-seq studies have captured only time points early in differentiation (up to two to four weeks^10,15,21,22^, or up to eight weeks^9,20,23^).

Furthermore, *in vitro* differentiation systems are prone to biological and technical variability influenced by replicates and differentiation protocols^24,25^, resulting in differences cell phenotype, as well as cardiac purity with the presence of other differentiated cell types in the population. Without extensive experimental and technical replicates in these types of transcriptomic studies, identification and validation of genes for more focused downstream analysis may be obscured or confounded by non-physiologically relevant factors. These issues have historically been challenging to address due to technical limitations in extended sample collection and storage, scRNA-seq experimental bottlenecks, and need for batch correction^26,27^.

To profile the dynamic cell populations during hiPSC differentiation to cardiomyocytes and their *in vitro* maturation after extended time in culture, we performed scRNA-seq on cells undergoing directed differentiation at Days 0, 12, 24, and 90. Differential expression analysis on de novo-identified cell clusters is often an important first step in scRNA-seq analysis to identify marker genes that distinguish cell types and cell states beyond the limited set of previously established markers^28,29^. We complemented this type of cluster-based differential expression analysis with a cluster-independent, time point-based penalized regression method^30,31^ that ranks and prioritizes genes based on how well they can predict the differentiation time point. The expression patterns of top-ranked genes were validated in re-plated cardiomyocytes by imaging using RNA Fluorescence in Situ Hybridization (RNA FISH). While some of the top-ranked genes were expected given their roles in cardiomyocyte development, this time point-based method allowed us to identify a short list of candidate genes that distinguished between D12, D24, and D90 cardiomyocytes with high accuracy. These data indicated that gene sets as small as two to twelve genes may be informative in staging cardiomyocyte maturation and differentiation state during extended culture.

In addition to characterizing transcriptional changes that occur during *in vitro* differentiation and maturation of cardiomyocytes, we explored the technical and biological variability arising from *in vitro* differentiation experiments. We sequenced cells from a total of 55 samples from multiple independent differentiation experiments with two endogenously fluorescently-tagged cells lines and their unedited parental line, each differentiated using two differentiation protocols. This reproducibility analysis was performed at the two earlier time points (D12, D24) with samples that were processed in a single batch to limit downstream batch effects. The differentiation experiments, cell lines, and protocols were correlated at the population level. However, we identified some variation in cardiomyocyte profiles within and across differentiation experiments. While fluorescent tags did not affect the transcriptional profiles of *in vitro* differentiated cardiomyocytes, we observed differences in the timing of key gene expression changes between cardiomyocytes based on differentiation protocol. Taken together, this work provides a comprehensive analysis of gene expression changes in a highly replicable and open-source dataset of differentiating cardiomyocytes.

## Results

### Single cell RNA-sequencing reveals distinct cell types and cardiomyocyte differentiation states after directed cardiomyocyte differentiation

To identify and characterize the distinct cell types and transcriptional states present during *in vitro* differentiation of hiPSC-derived cardiomyocytes, we performed scRNA-seq on cell populations spanning four stages of the cardiac differentiation process: undifferentiated hiPSCs (Day 0), early- and intermediate-stage cardiomyocytes (Day 12 and Day 24), and an aged cardiomyocyte population that served as a benchmark for more mature cardiomyocytes (Day 90)^32-34^. We used the scRNA-seq method SPLiT-Seq^35^ for parallel processing of all D12/D24 samples, including those from two differentiation protocols, three cell lines, and multiple independent differentiation experiments (sample overview shown in **Supplementary Fig. S1** and **Supplementary Table S1**).

We first focused our analysis on cells differentiated with a small molecule protocol (referred to as Protocol 1; **Fig. 1A, Supplementary Fig. S1B, D**) collected from all four time points (D0, D12, D24, D90; n = 11,619 cells), thereby establishing the baseline of cell types and gene expression patterns before expanding the analysis to the entire dataset. Unsupervised clustering identified 14 clusters representing three major categories of cells: undifferentiated hiPSCs, cardiomyocytes, and differentiated non-myocytes (**Fig. 1B**). The cluster corresponding to undifferentiated hiPSCs (C2) was identified by expression of the pluripotency transcription factor *POU5F1* (OCT-4) (**Fig. 1B-D**). Five of the 14 clusters contained cardiomyocytes based on the presence of the classic cardiomyocyte gene *TNNT2* (cardiac troponin T; C0, C1, C3, C7, C12; **Fig. 1C-D**), comprising the majority (∼72%) of the differentiated cell populations. Parallel measurement of cardiac muscle troponin T protein by flow cytometry in the same samples prior to sequencing confirmed the fraction of *TNNT2+* cells (R^2^ = 0.81; **Fig. 2B**). We observed cardiomyocytes with cell cycle activity (C12; co-expression of the proliferation marker *MKI67* and *TNNT2*) from all three differentiation time points (D12/D24/D90), with the number of *TNNT2*+ cells expressing *MKI67* decreasing with differentiation time point (n = 84 cells, 2.9% of D12 cardiomyocytes; 65 cells, 3.0% of D24 cardiomyocytes; and 42 cells, 2.0% of D90 cardiomyocytes; **Fig. 1D**). RNA FISH on re-plated cardiomyocytes was used to confirm the presence of *TNNT2*+/*MKI67*+ cardiomyocytes by imaging (**Fig. 1E**).

**Figure 1:**
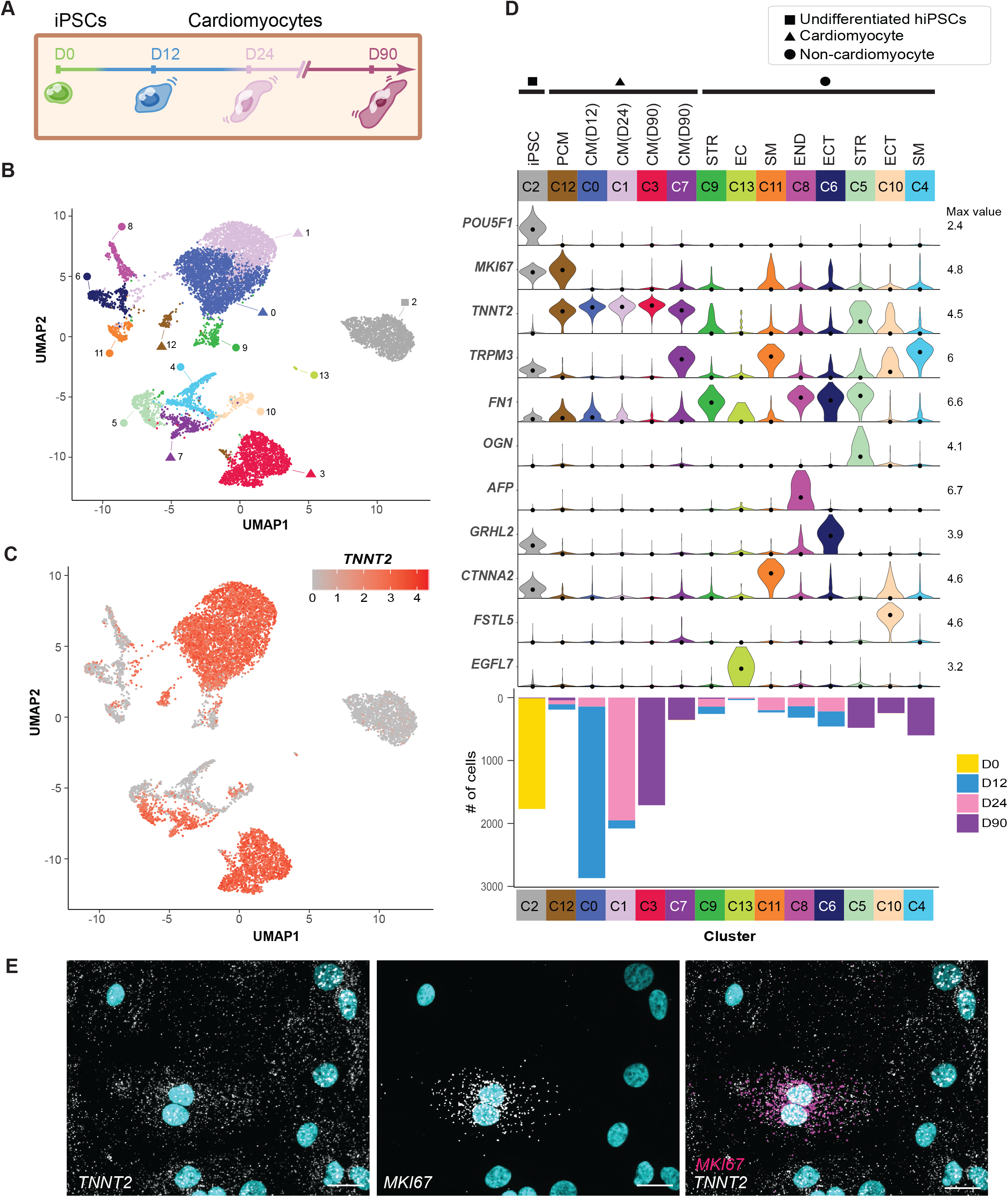
Unsupervised clustering reveals distinct cell types and cardiomyocyte clustering by time point. A. Sample collection schematic highlighting D0 hiPSCs and cell populations collected after 12, 24, and 90 days of cardiomyocyte differentiation with Protocol 1. B. Undifferentiated hiPSCs and hiPSC-derived cells present during cardiomyocyte (CM) differentiation collected at 3 time points (D0 = 1,764 cells, D12 = 3,538 cells, D24 = 2,854 cells, D90 = 3,463 cells; n = 11,619 total cells) were clustered using the Jaccard-Louvain method (14 clusters indicated by colors) and visualized using Uniform Manifold Approximation and Projection (UMAP). Cluster IDs were assigned after clustering based on cluster size, with C0 containing the most cells and C13 containing the least. Square icon identifies the cluster of undifferentiated hiPSCs (C2), triangles identify cardiomyocyte clusters (*TNNT2*+; C0, C1, C3, C7) including the proliferative cardiomyocyte cluster (*TNNT2*+/*MKI67*+; C12), and circles identify non-cardiomyocyte differentiated cell clusters (i.e., all other *TNNT2*-clusters). C. Same UMAP as B colored by transcript abundance of the cardiomyocyte marker cardiac troponin T (*TNNT2*). Of the differentiated cells (D12, D24, D90), 72% are *TNNT2+* cardiomyocytes. D. Violin plots showing normalized transcript abundance of cell type marker genes by cluster (max value = maximum value of log1p normalized counts; dot = median). Bar beneath each cluster indicates cluster size (# of cells) and is colored by time point. CM = cardiomyocyte, PCM = proliferative CM, STR = stromal-like, EC = endothelial-like, SM = smooth muscle-like, END = endodermal, ECT = ectodermal. E. Representative RNA FISH image showing *TNNT2* and *MKI67* transcripts in re-plated cardiomyocytes imaged at D30. Nuclei are labeled with DAPI (cyan). Scale bar = 20 µm

**Figure 2:**
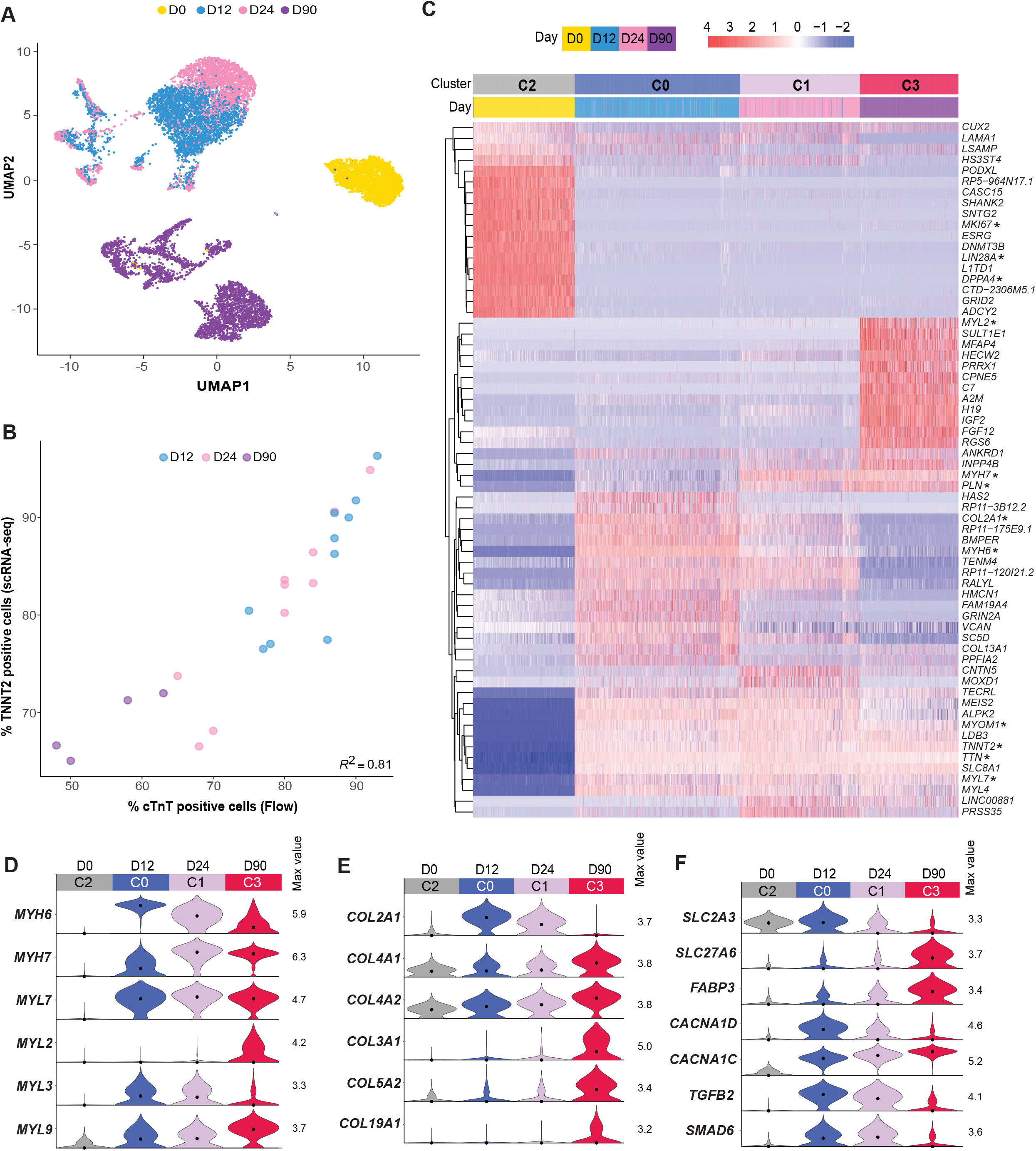
Transcriptional shifts between differentiation time points reveal changes in cardiomyocytes over time in culture. A. Same UMAP as shown in Fig. 1B, colored by collection time point; yellow -D0 undifferentiated hiPSCs; blue-D12 cells, pink-D24 cells, purple-D90 cells. B. Pearson correlation of cells expressing *TNNT2* by scRNA-seq and flow cytometry. The percent of TNNT*2*-positive cells in each sample from the scRNA-seq data is shown relative to the percent of cardiac troponin T (cTnT)-positive cells from each sample determined by flow cytometry. Individual points are colored according to time point; *R*^*2*^ = 0.81, *p* = 1.73e^-09^. C. Differentially expressed genes were identified between pairs of clusters corresponding to the undifferentiated hiPSC population (C2) and the three largest cardiomyocyte populations: D12 (C0), D24 (C1), and D90 (C3). Heatmap shows the top 10 up- and down-regulated genes from each pairwise cluster comparison. Normalized transcript abundance was centered and scaled across each row (z-score color scale on top; red = standard deviations above mean; blue = standard deviations below mean; white = mean; for visualization purposes, 4 was set as the maximum z-score, and z-scores > 4 were set to 4). The dendrogram is based on hierarchical clustering of genes. Each column corresponds to one cell. Genes referenced in Results are noted with an asterisk. D. Transcript abundance distributions for genes encoding myosin heavy and light chains with temporal transcriptional shifts between D0 (C2), D12 (C0) cardiomyocytes, D24 (C1) cardiomyocytes, and D90 (C3) cardiomyocytes. Max value = maximum value of log1p normalized counts; dot = median. E. Same as D but showing genes encoding collagens. F. Same as D and E, but showing genes encoding molecular transporters, ion channels, and signaling factors.

### Non-cardiomyocyte populations are distinct at the early and late time points

Consistent with previous studies of cardiac populations *in vitro*^9,10,36^ and the developing human heart *in vivo*^*6,37,38*^, we also observed non-cardiomyocytes (28% of cells across D12/D24/D90 by scRNA-seq) in the differentiated populations (**Fig. 1C, Fig. 2B, Supplementary Table S1**). In the two intermediate time points (D12/D24), non-cardiomyocytes were predominantly categorized into 4 clusters: *FN1+* stromal cells (C9) and smooth muscle-like cells expressing *TRPM3* and *CTNNA2* (C11), an endodermal subset expressing both *FN1* and *AFP* (C8), and an ectodermal cluster expressing *GRHL2* and *FN1* (C6; **Fig. 1D, Supplementary Fig. S2A-B, D-E**). There was also a small yet distinct cluster of endothelial cells, marked by the expression of *EGFL7* (C13; **Fig. 1D, Supplementary Fig. S2A)**. D90 non-cardiomyocytes were generally distinct from the D12 and D24 non-cardiomyocyte clusters, with C5, C10, and C4 indicative of stromal, ectodermal, and smooth muscle-like cells respectively (**Fig. 1D, Supplementary Fig. S2A**). Across all time points, the smooth muscle-like population (C11, D12/24; C4, D90) expressed high levels of *TRPM3*, with high expression of *CTNNA2* in the D12/24 time points (**Fig. 1D, Supplementary Fig. S2B**). Unlike at D12/D24, there was no dominant endodermal cluster in the late D90 population (C8, D12/D24; **Fig. 1D**).

In some cases, cell types were challenging to define due to the presence of multiple classical marker genes associated with cardiomyocytes and a secondary cell type within the same cluster. Specifically, C7 contained *TNNT2+* cardiomyocyte-like cells that also expressed *TRPM3* (**Fig. 1D, Supplementary Fig. S2D**), and C5 was comprised of cells expressing stromal (*FN1*) and ECM genes (*OGN*)^6,39^ with about half of these cells also expressing the cardiomyocyte marker *TNNT2* (**Fig. 1D, Supplementary Fig. S2C, E**).

### Gene expression changes during extended culture of immature cardiomyocytes are consistent with maturation over time

Immature cardiomyocytes continue to undergo gene expression changes after differentiation concurrent with the aging and maturation of cells *in vivo* and *in vitro*^6,40-43^. Within the cardiomyocyte (*TNNT2+*) populations, cells clustered along this temporal axis of differentiation with two dominant clusters of early/intermediate cardiomyocytes from D12/D24 (C0 and C1) and a distinct cluster of aged cardiomyocytes from D90 (C3; **Fig. 1B, Fig. 2A**). Transcriptional changes associated with these extended culture transitions were identified using standard pairwise differential expression comparisons between the four largest *TNNT2*+ clusters (C0-C3; **Fig. 1B, D, Fig. 2C**). As expected, there was a significant transcriptional shift between the undifferentiated hiPSC cluster (C2) and the three main cardiomyocyte clusters (C0, C1, C3), reflecting changes in cell proliferation (downregulated *MKI67*) and loss of stemness (downregulated *LIN28A, DPPA4*; **Fig. 2C**). Across all three differentiated cardiomyocyte timepoints, there was a broadly-expressed set of genes that was not temporally-regulated and included known cardiomyocyte structural genes *TNNT2* (cardiac troponin T), *TTN* (the sarcomere-spanning protein titin), and *MYOM1* (Myomesin-1; **Fig. 2C**). Consistent with the progression of genes associated with cardiomyocyte aging and maturation^11,44-46^, differentially expressed genes between early and intermediate stage cardiomyocytes (D12 and D24) related to the continuation of cardiac development and differentiation (i.e., gene ontology categories of “heart morphogenesis”, “regulation of membrane potential”, and “extracellular matrix organization”; **Supplementary Fig. S3C, Supplementary Table S4**). Between the intermediate and late stage cardiomyocytes (D24 to D90), enriched gene ontology categories included “cell-substrate adhesion” and “extracellular matrix organization”, in addition to “heart development” (**Supplementary Fig. S3C, Supplementary Table S4**).

Although measurably distinct from each other (**Fig. 1B, Fig. 2C**), the changes in gene expression between D12 and D24 were less pronounced than those seen between D24 and D90 with relatively modest shifts in median transcript levels between these two intermediate time points (**Fig. 2A, Fig. 2C-F, Supplementary Fig. S3C-D** and **Supplementary Table S2**). The myosin heavy chain genes *MYH6* and *MYH7* were among the relatively small set of genes with expression changes greater than two-fold from D12 to D24 (**Fig. 3A, Supplementary Table S2**). Across the population, *MYH6* was more abundant at D12, and *MYH7* was more abundant at D24 (**Fig. 2C-D**), indicative of an expression switch associated with the maturation of human ventricular cardiomyocytes^47,48^. C1 (mostly D24) also showed an increase in expression of the gene encoding cardiac calcium regulator phospholamban (*PLN*; **Fig. 2C**) and a decrease in expression of the gene encoding the voltage-dependent calcium channel Cav1.3 (*CACNA1D*) compared to cells in C0 (mostly D12; **Fig. 2F**).

**Figure 3.**
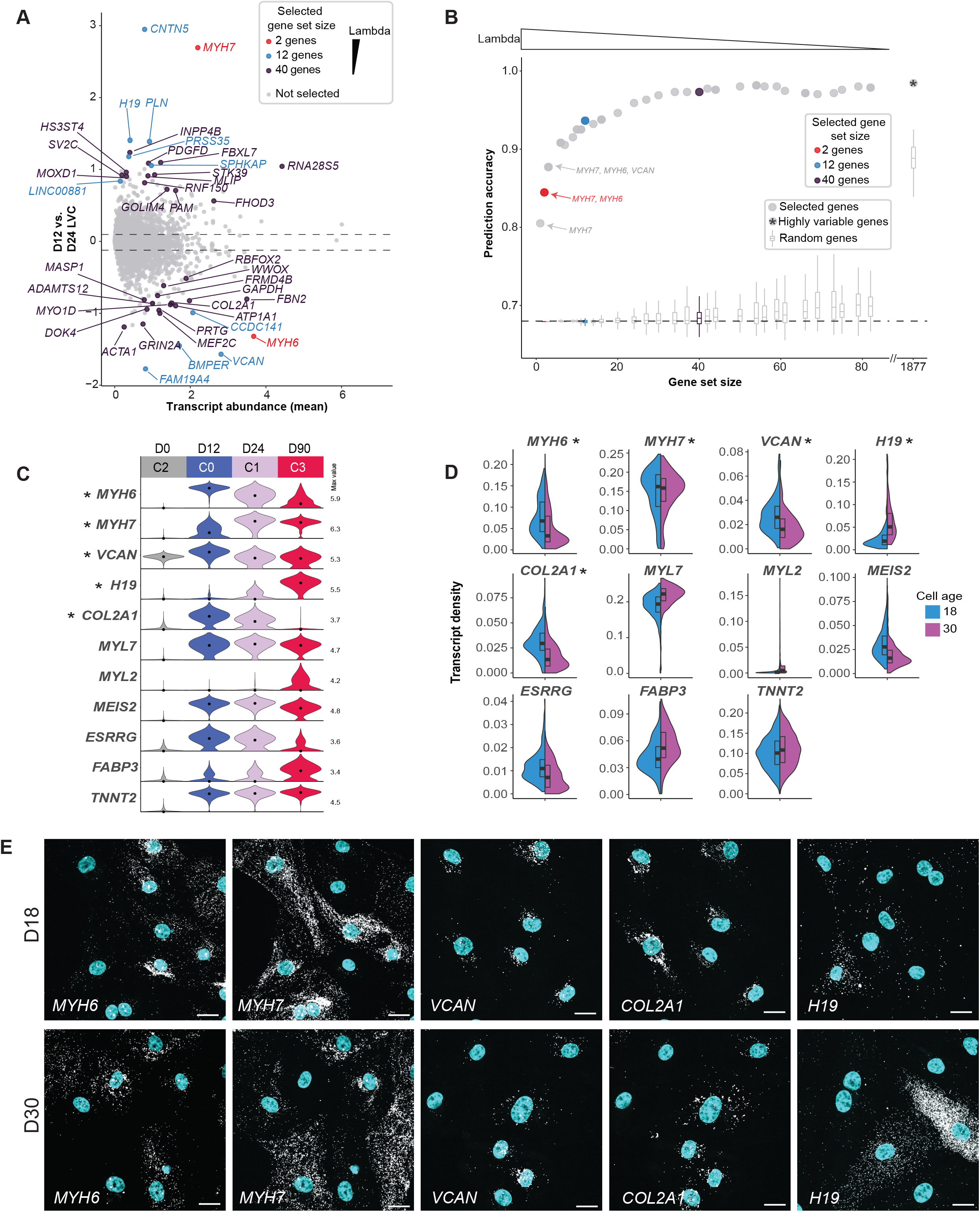
Bootstrapped sparse regression analysis identifies and ranks top differentially expressed genes for downstream analysis of D12 and D24 cardiomyocytes. A. Top 40 ranked genes identified as good predictors of cell age (D12 vs. D24) in bootstrapped sparse regression analysis with training data set are highlighted in a scatter plot of log2 fold change (LFC) between D12 and D24 vs. scRNA-seq mean transcript abundance (log1p of normalized counts; see Materials and Methods). Red, blue, and purple indicate gene sets selected at different values of the regularization parameter, lambda (red = 2 gene set at lambda = 0.487; blue = 12 gene set at lambda = 0.213; purple = 40 gene set at lambda 0.0494; selected gene sets are nested so that the 12 and 40 gene sets include the 2 and 12 gene sets, respectively). The remaining 23,625 genes that were not selected at any value of lambda are shown in gray. The dotted lines indicate 5% and 95% quantiles of log2 fold change. B. All unique selected gene sets (resulting from different values of the regularization parameter lambda) were used to calculate the prediction accuracy of cell age in the scRNA-seq bootstrapped sparse regression holdout data set (see Materials and Methods). X-axis shows each unique gene set size, and y-axis shows cell age prediction accuracy in holdout data. Color of selected gene set sizes are as follows: red = 2 gene set at lambda = 0.487; blue = 12 gene set at lambda = 0.213; purple = 40 gene set at lambda 0.0494; gray = all other gene set sizes. The prediction accuracy for a set of highly variable genes (n = 1,877) between D12 and D24 is shown as the dot to the right of the x-axis break. Prediction accuracies for random gene sets of the same size are shown as box plots with outliers omitted (selected gene sets ranged in size from 1 to 83 genes, and for each gene set size, 100 random gene samples were used for accuracy calculation). Dashed line at 0.68 indicates lower threshold for accuracy (holdout data set was 0.68 D12 and 0.32 D24 cells). C. scRNA-seq transcript abundance distributions for genes ranked in the top 40 in the D12 vs D24 bootstrapped sparse regression analysis (labeled with an “*”) and other genes of interest for downstream RNA FISH experiments. D. RNA FISH was performed on cardiomyocytes (images shown in panel E and Supplementary Fig. S5) that were replated onto glass at 12 days post-differentiation and allowed to recover for 5-6 days (D18, early time point; blue), or aged an additional 2 weeks (D30, intermediate time point; pink). RNA FISH transcript density (count/µm^2^) is shown for 11 genes (shown in panel C) in cardiomyocytes at the early and middle timepoint. Genes chosen from the bootstrapped sparse regression analysis are labeled with an “*”. Number of cells per probe target: *MYH6*: D18 = 418, D30 = 382; *MYH7*: D18 = 418, D30 = 382; *VCAN*: D18 = 169, D30 = 157; *H19*: D18 = 148, D30 = 117; *COL2A1*: D18 = 169, D30 = 157; *MYL7*: D18 = 539, D30 = 478; *MYL2*: D18 = 687, D30 = 595; *MEIS2*: D18 = 372, D30 = 283; *ESRRG*: D18 = 349, D30 = 291; *FABP3*: D18 = 148, D30 = 117; *TNNT2*: D18 = 405, D30 = 443. E. Representative images showing RNA FISH transcripts (white) for genes quantified in panel D with nuclei stained with DAPI (cyan). Representative fields of view are shown for both D18 and D30 time points. *MYH6* and *MYH7*; and *VCAN* and *COL2A1* were probed as pairs, and the same FOVs are shown for each pair. See Supplementary Fig. S5B for merged images of each pair. Images for additional genes are shown in **Supplementary Fig. S5A**. Scale bars = 20 µm.

The early/intermediate D12/D24 (C0, C1) to late D90 (C3) cardiomyocyte transition was marked by expression changes in cardiac muscle myosin genes often used to stage cardiac maturation, including the transition in expression from *MYH6* to *MYH7* and *MYL7* to *MYL2* in ventricular cardiomyocytes^49-51^ (**Fig. 2C, D**). In addition, we observed shifts from D24 to D90 in the ventricular and slow skeletal myosin light chain isoform *MYL3*, which decreased in expression over time in culture, as well as smooth muscle myosin light chain *MYL9*, which increased in expression over time in culture (**Fig. 2D**). Members of the SMAD family, which mediate TGF-beta signaling and are involved in early stage cardiac development and cardiomyocyte differentiation^52^, decreased in expression over time, while SMAD negative regulator *LDLRAD4* increased in expression during extended culture (**Fig. 2F, Supplementary Fig. S3D**). We observed other changes in expression in genes encoding components of signaling pathways including cyclic nucleotide (cAMP/cGMP) signaling (*PDE3A, PDE10A, PDE1C, ADCY5*, and ion channels (*CACNA1C, SCN5A* and *KCNQ1;* **Supplementary Fig. S3D**). Several collagens were upregulated in D90 cardiomyocytes (**Fig. 2E**), which is consistent with previous observations of collagen expression by cardiomyocytes during early *in vivo* heart development^11,53^. The collagen *COL2A1* decreased with age, a pattern previously reported to distinguish trabeculated cardiomyocytes from compact cardiomyocytes in human fetal heart development^6^ (**Fig. 2E**). Consistent with the known metabolic switch from glycolysis in hiPSCs to fatty acid oxidation in mature cardiomyocytes, D90 cardiomyocytes (C3) exhibited down-regulation of the glucose transporter *SLC2A3* and up-regulation of three metabolism associated genes: the long-chain fatty acid transport protein *SLC27A6*, the fatty acid binding protein *FABP3*^11^, and glucose response IGF-binding protein 5 (*IGFBP5*; **Fig. 2F, Supplementary Fig. S3D**).

### A complementary approach to identify subsets of differentially expressed genes for downstream analysis

Pairwise differential expression analysis (as shown above) is often used to identify genes of interest for downstream analysis such as functional validation or imaging^28^. This approach is cluster-dependent, with genes that differ between clusters identified at a pre-specified fold change or significance level (ex. Log2 fold > 1). We sought an alternative, complementary method to rank differentially-expressed genes and prioritize markers for downstream biological assays to reduce the number of genes that need to be tested experimentally. To accomplish this, we used a time point-based bootstrapped sparse regression statistical approach^30,31^ to identify and rank a subset of genes based on their ability to correctly assign individual cells to either the D12 or D24 cardiomyocyte time points in a training data set (see Materials and Methods; **Fig. 3A-B, Supplementary Fig. S4A-B, Supplementary Table S3**). Using only the expression level of the highest-ranked gene (*MYH7*) as input, a simple logistic model correctly assigned the cells in the holdout data set to D12 or D24 with an accuracy of 0.81 (**Fig. 3B**). Including the expression level of *MYH6*, the second ranked gene, improved the accuracy of prediction of time point for a given cell to 0.84 (**Fig. 3B – red dot**). The ranking of *MYH6* and *MYH7* as the top two genes is consistent with their changes in expression during early cardiomyocyte development^47,48^. Expanding the list to include the top 12 ranked genes raised prediction accuracy to 0.94 (**Fig. 3B – blue dot**). These included the cardiac calcium regulator phospholamban (*PLN*) as well as non-classical cardiac genes such as the proteoglycans *COL2A1* and *VCAN*. The top 40 genes resulted in a prediction accuracy of 0.97 (**Fig. 3B – purple dot**) and using 1,877 of the most highly variable genes between D12 and D24 cardiomyocytes led to a prediction accuracy of 0.98 (**Fig. 3B**). Thus, small numbers of genes can accurately classify differentiation time points, with diminishing returns by including additional genes in the model. One hundred percent of the top 12 ranked genes and 70% of the top 40 ranked genes were also identified as differentially expressed between C0 (D12) and C1 (D24) cardiomyocyte clusters, showing that the bootstrapped sparse regression method is consistent with standard cluster-based differential expression while providing additional ranking and prioritization of genes (**Supplementary Table S2, Supplementary Table S3**).

Of the top ranked genes at D12/D24, many showed a continued progression of up- or down-regulation between D24 and the D90 time point, indicating that their expression levels may be informative in identifying differentiation and maturation states at later time points (**Fig. 3C**). To further evaluate this, we repeated the bootstrapped sparse regression analysis on D24 and D90 cardiomyocytes and found similar prediction accuracy results with different genes (**Supplementary Fig. S4C-F**). The top three selected genes were able to assign cells to D24 or D90 with an accuracy of 0.94 and included *A2M*, a plasma protein secreted by cardiac fibroblasts, the maternally imprinted long non-coding RNA *H19*, and insulin-like growth factor *IGF2* (**Supplementary Fig. S4C, H, Supplementary Table S3**). The top 12 selected genes predicted the differentiation time point with over 0.98 accuracy, and the top 40 genes increased accuracy to 0.99 (**Supplementary Fig. S4D**). *H19* was the only gene shared in the top 12 of both the D12/24 and D24/90 analyses, and only four others, *COL2A1, BMPER, PRTG*, and *MYH6*, were shared in the top 40 genes across both D12/24 and D24/90 analyses (**Supplementary Fig. S3A-B, Supplementary Fig. S4C, Supplementary Table S3**). Overall, this feature selection analysis indicates that a small subset of genes contains most of the relevant information that discriminates between the transcriptional profiles of D12/D24 and D24/D90 cardiomyocyte populations.

### Validation of select genes using RNA FISH

We used RNA FISH to validate the expression patterns of select genes by quantifying transcripts in cardiomyocytes re-plated on glass for high-resolution imaging. Due to the technical challenges of culturing single cells out to D90, we performed RNA FISH only at the early and intermediate time points (**Fig. 3E, Supplementary Fig. S5**; cells were re-plated at D12 and allowed to recover for six days, or aged an additional 18 days for D18/D30 time points; see Materials and Methods). We targeted 11 genes for follow-up by RNA FISH to confirm expression in re-plated cardiomyocytes, with five genes chosen from the D12/24 top 40 bootstrapped sparse regression list: *MYH6, MYH7, VCAN, COL2A1*, and *H19*. The remaining genes were chosen for being differentially expressed between cardiomyocyte clusters (C0/D12, C1/D24, C3/D90) and for their known roles in cardiomyocyte development (**Fig. 3C**). Transcript abundance was quantified in segmented cells at each time point, and most assayed genes showed trends consistent with the scRNA-seq data (**Fig. 3C-E**). Of the top two performing genes predictive of D12/D24 in the bootstrapped sparse regression analysis (*MYH6* and *MYH7*), RNA FISH showed an expression change for *MYH6*, which decreased in expression between D18 and D30 (**Fig. 3D**). *MYH7* did not show a large expression change with RNA FISH. While the sample time points are similar, re-plating cells for RNA FISH delays the time points relative to scRNA-seq (D18/30 vs D12/24) and places the cells on a stiffer substrate^50^. Furthermore, while *MYH6* continued to decrease after D24 in the scRNA-seq data (**Fig. 3C**) and was among the top ranked genes between D24 and D90, *MYH7* was not a top ranked gene in the bootstrapped sparse regression analysis between D24 and D90 (**Supplementary Fig. S4C, Supplementary Table S3**). This is consistent with the switch from *MYH6* to *MYH7* expression occurring before D18 and the observed stable expression of *MYH7* between the D18 and D30 RNA FISH time points. The scRNA-seq analysis revealed a range of *MYH6* and *MYH7* expression that was largely anti-correlated in single cells, and this cell-to-cell heterogeneity was also confirmed by RNA FISH (**Supplementary Fig. S4G, Fig. 3D**). Finally, the RNA FISH confirmed the expression of non-classical cardiac genes such as *H19* and *COL2A1* in differentiated and re-plated cardiomyocytes (**Fig. 3E**).

### Expanding early and intermediate time point analysis to explore reproducibility of differentiation

Analysis of differentiated cell populations from Protocol 1 revealed distinct cell types and changes in gene expression over the time course of differentiation and established a baseline for applying these interpretations to a broader dataset. *In vitro* differentiation systems are prone to biological and technical variability, potentially arising from variables including hiPSC line, culture conditions, and differentiation protocol steps. To assess the robustness of our protocols and analysis methods, we expanded our data set to characterize the reproducibility of cardiomyocyte differentiation in different protocols and hiPSC clones. We sequenced samples from the two intermediate time points that were differentiated with a second differentiation protocol that used a combination of cytokines and small molecules (Protocol 2). While cells for both protocols were collected on the same day, the nomenclature reflects a two day difference (i.e., D12 for protocol 1 is the same as D14 for protocol 2, D24 for Protocol 2 is the same as D26 for protocol 2; see Materials and Methods). We also collected samples from multiple cell lines in the WTC-11 background and independent differentiation experiments. This resulted in a total of 15,878 D12/14/24/26 sequenced cells (n = 7,987 cells in Protocol 1, n = 7,891 cells in Protocol 2), representing 48 independent samples (**Fig. 4A-D, Supplementary Fig. S1C, E;** seven samples from D0/D90 are not included here). To limit downstream batch effects that might affect this comparison, all samples from intermediate differentiation time points were processed in a single library preparation and sequencing batch.

**Figure 4.**
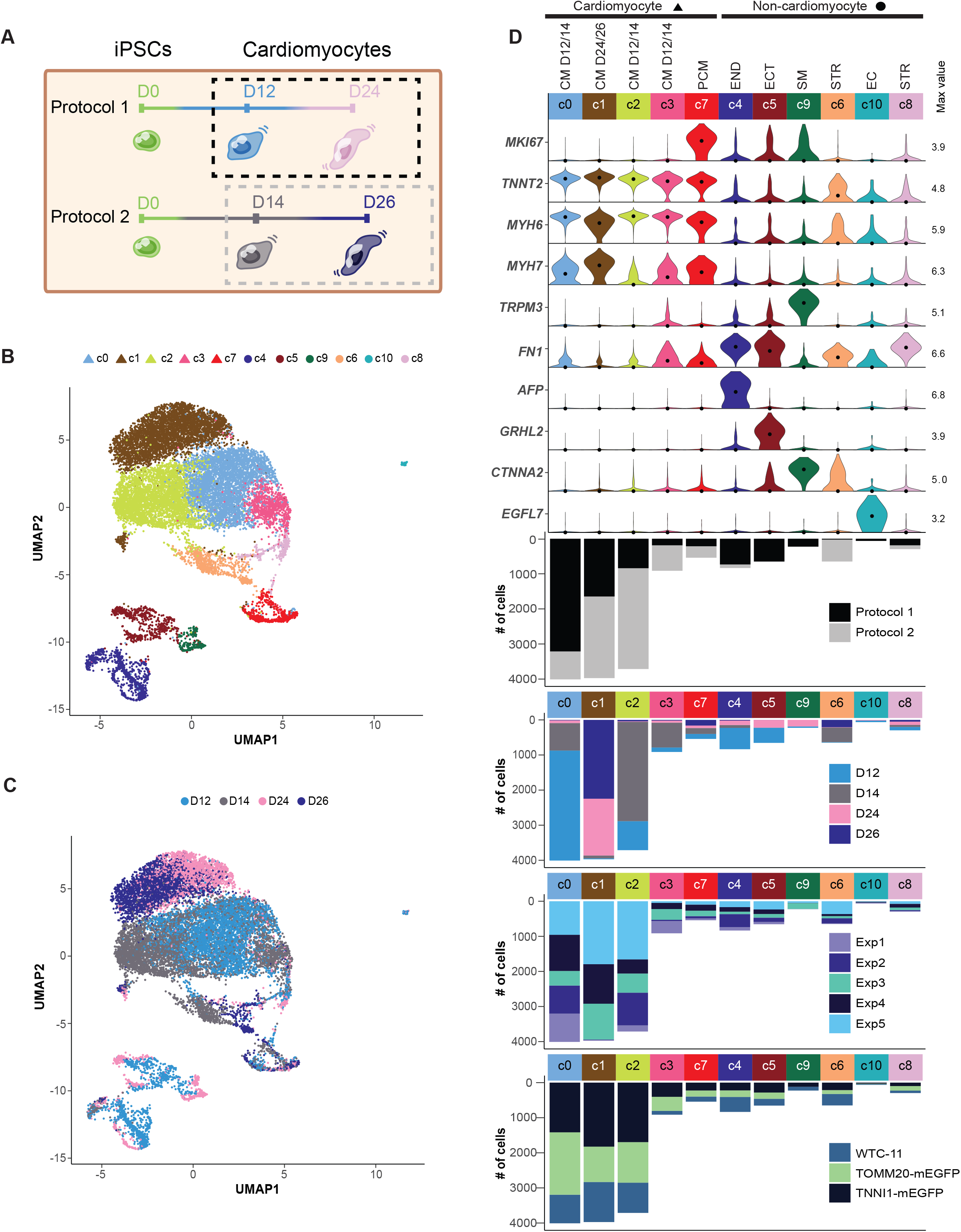
Expanded analysis of samples collected 12, 14, 24, and 26 days after the initiation of differentiation with two directed differentiation protocols. A. Sample collection schematic highlighting collection of cells at early and intermediate time points (dashed boxes) from Protocol 1 (small molecule, D12/24 cells) and Protocol 2 (cytokine, D14/26 cells). B. UMAP of early and intermediate time point cells differentiated from both protocols, representing three clonal cell lines and five independent differentiation experiments, Cells are colored by cluster (11 clusters), and shape indicates cardiomyocyte vs non-cardiomyocyte cluster (triangle = *TNNT2*+ cardiomyocytes, circle = non-cardiomyocytes). Number of cells: 5,494 D12 cells, 5,135 D14 cells, 2,493 D24 cells, 2,756 D26 cells. C. Same UMAP shown in panel B, colored by collection time point; Protocol 1 (small molecule) = D12 and D24. Protocol 2 (cytokine) = D14 and D26. D. Violin plots showing distributions of transcript abundance for cell type marker genes by cluster from panel B (max value = maximum value of transcript abundance; dot = median). Bar beneath each cluster indicates cluster size (# of cells) and is colored by protocol, time point, differentiation experiment, and cell line.

We found that D12/D14/D24/D26 cells clustered into four non-proliferative cardiomyocyte clusters (c0, c1, c2, c3; *TNNT2+*, clusters in this analysis are distinguished with a lowercase “c” in **Figs. 4, 5**, and **Supplementary Figs. S6, S7**), one proliferative cardiomyocyte cluster (c7; *TNNT2+*/*MKI67*+), and six *TNNT2-*non-cardiomyocytes clusters (c4, c5, c6, c8, c9, c10) (**Fig. 4B-D**). c1 was almost exclusively composed of D24 and D26 cells and had the highest median *MYH7* level and lowest median *MYH6* level among the cardiomyocyte clusters (**Fig. 4D**), consistent with changes in expression of *MYH6* and *MYH7* during ventricular cardiomyocyte development. The other three non-proliferative cardiomyocyte clusters (c0, c2, c3) were dominated by D12/D14 cells and differed in their median level of *MYH7* expression with c2 being the lowest and c0 being the highest (**Fig. 4D**).

**Figure 5.**
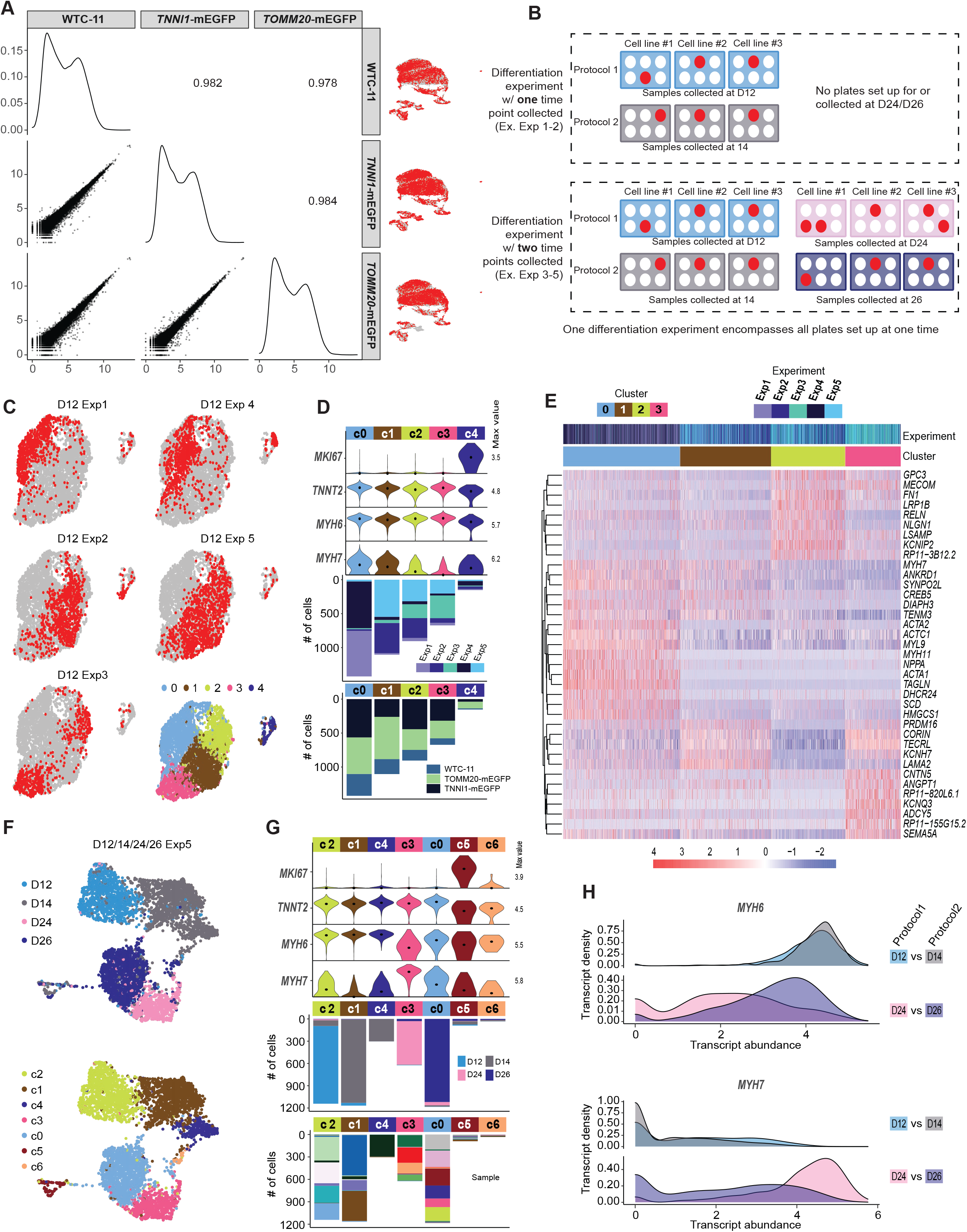
Similarities and differences in gene expression between cell lines, differentiation experiments, and differentiation protocols. A. Scatter plots of population transcript abundances between three different cell lines: two clonal gene-edited lines with endogenous fluorescent tags: AICS-0011 TOMM20-mEGFP and AICS-0037 TNNI1-mEGFP, and their parental line WTC-11. Each point represents a gene. Spearman correlations are shown in upper right. UMAPs on right highlight cells from each cell line in red. B. Differentiation experiment schematic. One differentiation experiment includes all plates set up on that experiment date and may include multiple cell lines, protocols, and/or time points. One plate or one half-plate was set up for each intended collection time point per cell line and protocol. Differentiation experiments were seeded on different days using an independent source plate of hiPSCs. Each scRNA-seq sample originates from a single well in a differentiation plate; in some cases, multiple wells/samples were collected per plate but were never pooled. C. Cardiomyocytes (*TNNT2*+ cells) from all collected D12 samples (from each of the five differentiation experiments) were independently clustered and visualized using UMAP. Lower right UMAP is colored by cluster, and each other UMAP highlights in red cells from one of the five differentiation experiments. D. Group violin plot showing distributions of marker genes in D12 clusters with cluster breakdown by experiment and cell line shown in the bar plots below. E. Heat map of top differentially expressed genes between the four non-proliferative (*MKI67-*) D12 cardiomyocyte clusters (c0, c1, c2, c3). Normalized transcript abundance was centered and scaled across each row (z-score color scale below heatmap; red = standard deviations above mean; blue = standard deviations below mean; white = mean; for visualization purposes, 4 was set as the maximum z-score, and z-scores > 4 were set to 4). The dendrogram is based on hierarchical clustering of genes. Each column corresponds to one cell. F. Experiment 5 cardiomyocytes were independently clustered to compare differentiation protocols. Top: UMAP colored by day of differentiation (Protocol 1 = D12, D24; Protocol 2 = D14, D26). Bottom: UMAP colored by cluster. G. Group violin plot showing distributions of marker genes in Experiment 5 cardiomyocyte clusters with cluster breakdown by day and by sample shown below. H. Distributions of *MYH6* (top) and *MYH7* (bottom) transcript abundance in differentiation experiment 5. Protocol 1 = D12, D24; Protocol 2 = D14, D26.

### Contribution of cell line and experimental replicates to variability in differentiation

We calculated gene-wise coefficients of determination (R^2^) for variables of interest (time point, cell line, protocol, differentiation experiment) to quantify sources of variation and found that time point was the major source of variation as expected. While differentiation protocol and differentiation experiment also contributed to variance in gene expression, the specific cell line was not a major contributor (**Fig. 5A, Supplementary Fig. S6A**). The D12/D14 and D24/D26 samples included three hiPSC lines in the same genetic background (two clonal gene-edited lines with endogenous fluorescent tags: AICS-0011 TOMM20-mEGFP and AICS-0037 TNNI1-mEGFP, and their parental line WTC-11). Expression profiles across the three cell lines were highly correlated at the population level, and cells did not cluster by cell line, indicating that the presence of a fluorescent tag on *TOMM20* or *TNNI1* genes and the clonal selection process used to generate these lines did not alter the differentiation potential of hiPSCs or the transcriptional profiles of differentiated tagged cardiomyocytes (**Fig. 4D, Fig. 5A**).

Gene expression in samples derived from independent differentiations were also correlated at the population level (**Supplementary Fig. S6B**). To explore experimental differences that may be obscured in the global analysis, we independently clustered cardiomyocytes from each time point (Clusters c0, c1, c2, c3, and c7 in **Fig. 4B-D**) and evaluated the extent to which cells clustered by experiment **(Fig. 5C-D, Supplementary Fig. S6C-E**; see **Fig. 5B** for definition of differentiation experiment). Overall, most clusters contained cells from a mix of multiple differentiation experiments, however a few clusters were dominated by one or two differentiation experiments at each time point (D12 time point is shown in **Fig. 5C, D**, all others in **Supplementary Fig. S6C-E**). These clusters did not appear to represent distinct cardiomyocyte cell types, but rather committed cardiomyocytes at different stages along the cardiomyocyte maturation trajectory. This conclusion was supported by differences in the ratio of expression of *MYH6* to *MYH7* between clusters (**Fig. 5D, Supplementary Fig. S6C-E**). Additional differentially expressed genes within time points included smooth muscle genes (*TAGLN* and *MYL9*) and early atrial genes (*NPPA* and *ACTC1*), both of which tended to be higher in the Exp1 and Exp4 populations at D12 (c0) compared to Exp2, Exp3, and Exp5 at D12 (**Fig. 5D, E**). Despite these differences, cardiac troponin T levels, as detected by flow cytometry and by *TNNT2+* expression, were similar across these samples, suggesting that cardiac troponin T alone may not sufficiently capture the biological variation across independent differentiation experiments and that more refined descriptors of cardiomyocyte state are needed (**Fig. 5D, Supplementary Fig. S6C-E, Supplementary Table S1**).

### Cell types and gene expression are generally reproducible across two differentiation protocols with variable timing of the *MYH6/MYH7* expression switch

Finally, we compared the composition of cell populations and gene expression between samples collected from two cardiomyocyte differentiation protocols. At the population level, gene expression profiles were well-correlated for the two protocols at both the early (D12/D14) and middle (D24/D26) collection time points (r*S* = 0.97 at D12/14, and r*S* = 0.96 at D24/26; **Supplementary Fig. S7A**). While both protocols showed similar *TNNT2* transcript levels (**Fig. 4D, Fig. 5G, Supplementary Fig. S7C-F**), Protocol 1 generated more non-cardiomyocytes in our experiments (24% cells for protocol 1 vs. 11% cells for Protocol 2: **Fig. 4D**, black vs gray bars). Some of the non-cardiomyocyte clusters were protocol-specific, with smooth muscle (c9 - Protocol 1, *CTNNA2* positive), ectodermal cells (c5 - Protocol 1, *GRHL2* positive), endodermal cells (c4; protocol 1, *AFP* positive), and endothelial cells (c10; protocol 1, *EGFL7* positive; **Fig. 4D**) all being more prevalent in Protocol 1. The stromal population (c8, both protocols, *FN1* positive) was generated by both protocols (**Fig. 4D**).

We next independently re-clustered cardiomyocytes from the largest differentiation experiment, Exp5 (**Fig. 5F, G**), to focus on protocol-specific differences within a high-replicate experimental setup. We found that cells clustered by differentiation protocol, and at the D24/D26 time point, there was a distinct trend for higher levels of *MYH7* and lower levels of *MYH6* in Protocol 1 compared to Protocol 2 (**Fig. 5G-H**). This trend of reduced *MYH7* expression in Protocol 2 was also found for the earlier time point (D12/D14) and was observed in three of the other independent experiments (**Supplementary Fig. S7C-F**). While these results do not indicate that the two protocols generate distinct cardiomyocyte cell types, they suggest that the timing of the *MYH6*/*MYH7* switch may be slightly different between the protocols within the first few weeks of differentiation. This difference may result from different inductive factors provided to the differentiating cells in the two protocols or from the presence of different types and numbers of non-cardiomyocytes in the differentiating populations. To further compare the gene expression changes in cardiomyocytes generated from both protocols, we repeated the bootstrapped sparse regression analysis to identify genes that best discriminated between D14/D26 cardiomyocytes derived from Protocol 2. We found that many of the top genes overlapped with those identified in the analysis of Protocol 1 (*MYH6, MYH7, VCAN, BMPER, CTNT5, H19, COL2A1, PLN, PRSS35, MEF2C*; **Supplementary Fig. S7B, Supplementary Table S3**), many of which were also among the top differentially-expressed genes between all four early/intermediate cardiomyocyte time points (D12/D14/D24/D26, **Fig. 2C, Supplementary Fig. S3A, Supplementary Fig. S7G**). These data suggest that the cells from both protocols undergo a similar progression of differentiation and maturation in these first few weeks and that there is a subset of genes that can robustly distinguish between time points across both differentiation protocols.

## Discussion

In this study, we used single cell RNA-sequencing to profile the cell populations present during hiPSC differentiation to cardiomyocytes and *in vitro* maturation after extended time in culture. Analysis of the early and intermediate time points revealed cell types and transitions consistent with previously reported studies, with cardiomyocytes clustering predominantly by time point and the presence of distinct non-cardiomyocyte clusters^10,23^. Extending the sample collection to 90 days post-differentiation provided insight into changes in population composition and gene expression at the single cell level in more mature cardiomyocytes. The transition between D24 and D90 cardiomyocytes encompassed changes in many of the structural, metabolic, and signaling transcriptional programs that are activated during *in vivo* cardiomyocyte development^9^. We identified several clusters that suggest *in vitro* cardiomyocyte maturation. Interestingly, we observed a subpopulation of D90 cardiomyocytes with upregulated expression of collagens and extracellular matrix-associated genes (C5, **Supplementary Fig. S2A**). This observation is consistent with previous findings of collagen expression during early *in vivo* heart development, perhaps indicative of a transient subpopulation of cardiomyocytes with extracellular-matrix related gene expression in the heart during early cardiac development^11,53^, but could also be a technical artifact resulting from doublets produced during sample processing. At D90, we observed multiple clusters displaying co-expression of classical cardiomyocyte genes and another secondary cell type, including a mixed cardiomyocyte-like population expressing the cardiac smooth-muscle TRP channel gene *TRPM3* (C7, **Fig. 1D, Supplementary Fig. S2A, D**). The TRPM3 channel is activated by thermal and hypotonic conditions^38^, thus we postulate that co-expression in this context may be a consequence of the *in vitro* culture conditions.

*In vitro* cardiomyocyte differentiation is prone to biological and technical variability that may be driven by factors such as the genetic background of cell lines, the batch of cells and other reagents used at the outset of an experiment, cell cycle and density of undifferentiated hiPSCs, and the protocols used for cell culture and directed differentiation^9,54-58^. This variability is a challenge for data reproducibility and makes it difficult to draw conclusions about cell types or states based on a limited number of samples and conditions. Historically, evaluation of cardiomyocyte differentiation performance and quality control has been evaluated using a single metric: expression of cardiac troponin T by flow cytometry or immunochemistry^56^. However, differentiated populations vary not only in the percent of cells expressing cardiac troponin T, but also exhibit variation in functional phenotypes including contractility, sarcomere organization, and electrophysical properties^56,57,59^.To probe the robustness of transcriptional profiles at the single cell level, we multiplexed the single cell analysis of >15,000 cells from 48 independent samples, spanning two differentiation protocols, three edited cell lines, and numerous experimental replicates. We found that while population level gene expression was well-correlated across experimental replicates, cell lines, and protocols, there were some differences that were not fully captured by pre-sequencing analysis of cardiac troponin T by flow cytometry. This heterogeneity included expression differences in *MYH6* and *MYH7* across experimental replicates and differences in genes associated with atrial or ventricular specification and smooth muscle (**Fig. 5C-H, Supplementary Figs. S6C-E, S7C-F**).

Transcriptional differences in cardiomyocytes between differentiation time points were evaluated using pairwise, cluster-based differential expression analysis, a common approach in analyzing scRNA-seq data. We complemented this analysis with a time point based, cluster-independent penalized regression method, which enabled ranking and prioritization of genes based on how well they predicted the differentiation time point. While some of the top genes identified were of known importance in cardiomyocyte biology, we observed a number of genes that have little to no previously reported role in cardiac differentiation or maturation. For example, the top three genes in the D24/D90 analysis were a plasma protein secreted by cardiac fibroblasts (*A2M*), long non-coding RNA *H19*, and insulin-like growth factor (*IGF2)*. While the collective role of these three genes in cardiac maturation has not been robustly established, *H19* and *IGF2* are co-expressed during development^60-62^. Furthermore, *H19* has been found to inhibit the abundance of the cardiac maturation-inducing micro-RNA let7^63^, and *A2M* has been reported to promote cardiomyocyte hypertrophy in ventricular cardiomyocytes^64^. Their performance in this prediction model suggests they may be important for distinguishing cardiomyocyte maturation states in other studies. Notably, this analysis revealed that gene sets as small as two to twelve genes enabled prediction of time point with high accuracy, similar to the accuracy achieved by using over 1000 of the most highly variable genes. These data indicate that a small subset of carefully-chosen gene targets may be informative for downstream studies where gene set size is limited, such as in functional knock-out assays, *in vivo* experiments, or image-based RNA FISH studies^65^.

In summary, we used scRNA-seq to profile gene expression in cardiomyocytes and non-cardiomyocytes during cardiomyocyte differentiation and extended culture *in vitro*. We tested the robustness of our conclusions by sequencing 55 total samples from numerous differentiation experiments, differentiation protocols, and cell lines. We found that while cell types and gene expression were generally correlated at the population level, there were differences in cardiomyocyte gene expression by differentiation protocol and experimental replicate. Using a cluster-independent regression analysis, we identified sets of two to forty genes that predict cardiomyocyte time point with high accuracy. This shows that a limited number of genes can be used to benchmark the stage of cardiomyocyte differentiation and maturation, providing insight into the quality of cardiomyocytes for use in subsequent *in vitro* models or *in vivo* therapeutic applications.

## Acknowledgements

We thank Natalie Gaudreault for imaging assistance, Derek Thirstrup for Cell Profiler analysis, Melissa Hendershott for cell imaging advice, and Haseeb Malik for cell maintenance during RNA FISH studies. We thank Thao Do for illustrations, Susanne Rafelski for thoughtful discussions on imaging and quantitation, and Gokhan Dalgin and Natalie DeWitt for editing assistance on the manuscript text. We thank the Allen Institute for Cell Science team for helpful scientific discussions and support. The WTC-11 line that we used to create our gene-edited cell lines was provided by the Bruce R. Conklin Laboratory at the Gladstone Institute and UCSF. We gratefully acknowledge funding from R21CA246358 and support from the Washington Research Foundation. We would like to thank the Allen Institute for Cell Science founder, Paul G. Allen, for his vision, encouragement, and support.

## Declaration of Interests

A.B.R, C.R. and G.S. are shareholders of Parse Bioscience.

**Please refer to Supplementary Materials for Supplementary Figures, Legends, and Tables**.

## Methods

### Human induced pluripotent stem cell culture

All human induced pluripotent stem cell (hiPSC) work was approved by internal oversight committees and conducted in accordance with NIH, NAS, and ISSCR guidelines. Cell lines used in this manuscript were: AICS-00 (WTC-11 unedited parental line, generously provided by Dr. Bruce R. Conklin, The Gladstone Institute^66^); and edited lines generated as described previously^67,68^: AICS-0011 cl.27 (TOMM20-mEGFP), AICS-0037 cl.172 (TNNI1-mEGFP). Edited cell lines can be obtained through the Allen Cell Collection (www.allencell.org/cell-catalog), and the unedited WTC-11 parental line can be obtained from Coriell (GM25256). Undifferentiated hiPSC lines were maintained on plates coated with growth-factor-reduced matrigel (Corning #354230) in mTeSR1 (Stem Cell Technologies #85850) supplemented with 1% penicillin/streptomycin (P/S) (Gibco #15070063). Cells were passaged after reaching 70-85% confluency (every 3-4 days) using Accutase (Gibco #A11105-01) to disperse into single cells and replated in mTeSR1 supplemented with 1% P/S containing 10 µM Rock Inhibitor (Y-27632, Stem Cell Technologies #72308). A full protocol is available at the Allen Cell Explorer (www.allencell.org/methods-for-cells-in-the-lab SOP: WTC culture v1.7.pdf).

### Cardiomyocyte differentiation using two protocols

Two differentiation protocols were used, referred to as “Protocol 1” and “Protocol 2” in the text. Protocol 1 is based on a previously reported small molecule differentiation method with modifications^69,70^. HiPSC cultures were dissociated into single cells using Accutase and plated into matrigel-coated 6 well-plates at 125k - 350k cells per well in mTeSR1 supplemented with 1% P/S and 10 µM Rock Inhibitor (Day -3). Media was replaced for 2 days, and differentiation was initiated on the third day (Day 0) by replacing media with RPMI-1640 (Invitrogen #A10491-01) supplemented with B27 without insulin (Invitrogen #A1895601) and 6 µM CHIR99021 (Cayman Chemical #13122). Media was replaced after 48 hours (Day 2) with RPMI-1640 supplemented with B27 without insulin and 5 µM IWP2 (R&D Systems #3533). Media was again replaced after an additional 48 hours (Day 4) by RPMI-1640 supplemented with B27 without insulin. Every 2-3 days thereafter (starting on Day 6), media was replaced with RPMI-1640 supplemented with B27 containing insulin (Invitrogen #12587010) and 1% P/S. Cardiomyocyte samples at Day 90 were differentiated using an optimized version of Protocol 1 with Chiron and IWP2, both at 7.5 µM. A full protocol is available at the Allen Cell Explorer (www.allencell.org/methods-for-cells-in-the-lab,SOP: Cardiomyocyte differentiation methods_v1.2.pdf).

Protocol 2 is a previously reported method using a combination of cytokines and small molecules to induce cardiac differentiation^71^. Briefly, hiPSCs were dissociated into a single cell suspension using Accutase and seeded into matrigel-coated 6 well-plates at 1×10^6^ -2×10^6^ cells per well in mTeSR1 supplemented with 1% P/S, 10 µM Rock inhibitor, and 1 µM CHIR99021 (denoted as Day -1). Differentiation was initiated the following day (Day 0) with the addition of RPMI-1640 supplemented with B27 without insulin, 100 ng/mL ActivinA (R&D Systems #338-AC), and 1:60 diluted growth-factor-reduced Matrigel. After 17 hours (Day 1), media was replaced with RPMI supplemented with B27 without insulin and containing 1 µM CHIR99021 and 5 ng/ml BMP4 (R&D systems #314-BP). After an additional 48 hours (Day 3), media was replaced with RPMI-1640 supplemented with B27 without insulin and 1 µM XAV 939 (Tocris Biosciences #3748). After an additional 48 hours (Day 5), media was again replaced with RPMI-1640 supplemented with B27 without insulin, and cultures were fed with RPMI-1640 supplemented with B27 containing insulin and 1% P/S every 2–3 days thereafter (starting on Day 7).

Across both differentiation protocols, spontaneous beating was generally observed between Days 7-12. We have recently updated our protocols and recommend the use of RPMI-1640 (Invitrogen #11875-093) and B27 supplement (Gibco #17504044) for cardiac differentiation using both protocols.

### Experimental design for scRNA-seq studies

Each differentiation experiment encompassed all plates set up at one experiment date, and one source plate of hiPSCs was dissociated and seeded concurrently for both differentiation protocols, thus keeping the input hiPSCs the same within an experiment. Differentiation experiments were seeded on different days using an independent source plate of hiPSCs. Samples were collected on the same day for each paired collection time point listed in the scRNA-seq dataset, from both protocols (Day 12/D14, D24/D26). The difference in reported day (i.e., Day 12 vs 14) is due to the delay in differentiation initiation (Day 0) between the two protocols, as described above in the “Cardiomyocyte differentiation using two protocols” section above. Samples were collected 15 days after seeding (denoted as D12 for Protocol 1, D14 for Protocol 2) or 27 days later (denoted as D24 for Protocol 1, D26 for Protocol 2). Samples referred to as D90 were independently derived and were collected 93-96 days after initiating differentiation using a modified version of Protocol 1 as described above. Each scRNA-seq sample originates from a single well in a differentiation plate. In some cases, multiple wells/samples were collected per plate but were never pooled. Stem cell (D0) samples for scRNA-seq were independently cultured and were not used as a source plate for any of the differentiation setups. There was a total of 55 samples included in this study; all scRNA-seq sample metadata can be found in **Supplementary Table S1**.

### Single cell dissociation for scRNA-seq

Stem cells were dissociated to single cells as described above and processed for RNA sequencing following the protocol detailed in the “scRNA-seq library preparation and sequencing” section below. All differentiated sample wells were visually inspected at the desired cardiomyocyte collection time point for successful cardiac differentiation on the basis of high cardiac purity, spontaneous beating, and morphology; wells that passed these criteria were collected for downstream analysis. For single cell dissociation, cardiomyocyte wells were washed with 1X DPBS (Gibco #14190-144) and incubated with pre-warmed 2X TrypLE Select (Gibco #A12177) diluted in Versene (Gibco #15040-066) for 8-10 minutes at 37°C. Monolayers were gently dissociated using a P1000 micropipette to obtain single cells, collected, and added to 5 mL of resuspension media - RPMI-1640 containing B27 supplement, 1% P/S, 10 µM Rock Inhibitor and 200 U/mL DNAse I (Millipore Sigma #260913-10MU). Wells were washed twice with an additional 1 mL of resuspension media, and the cell suspension was centrifuged at 300 g for 5 minutes at 4°C. The single cell suspension was gently resuspended in 5 mL of resuspension media and counted twice on a hemocytometer to obtain a total cell count for each sample.

### Flow cytometry of scRNA-seq samples

An aliquot was taken from each cell sample at the time of single cell dissociation for cardiac Troponin T (cTnT) analysis by flow cytometry. Briefly, sample aliquots were fixed with 4% paraformaldehyde (Electron Microscopy Sciences #15710) in DPBS for 10 min. After fixation, samples were stained for 30 min in BD Perm/Wash™ buffer (BD Biosciences #512091KZ) with anti-cardiac Troponin T AlexaFluor® 647 (BD Biosciences #565744) or an equal mass of AF647 lgG1 κ isotype control (BD Biosciences #565571). Fixed cells were resuspended in 5% FBS (Gibco #10437028) in DPBS with 2 µg/mL DAPI, followed by processing and data acquisition on a CytoFLEX S (Beckman Coulter). Analysis was performed using FlowJo software V. 10.2 (Treestar).

### scRNA-seq library preparation and sequencing

ScRNA-seq was performed using the SPLiT-seq method^35^. After single cell dissociation (see “Single cell dissociation for scRNA-seq”) samples were prepared for library preparation by centrifuging the cell suspension at 300 g for 5 minutes at 4°C and then resuspending in 1 mL of cold RNAse-free PBS containing 0.05 U/µL Superase IN (Invitrogen #AM2696) and 0.05 U/µL Enzymatics RNAse Inhibitor (Qiagen #Y9240L) (mixture referred to throughout as PBS + RI). On ice, resuspended cells were passed through a 40 µm filter and fixed with 3mL of 1.33% formaldehyde (Electron Microscopy Sciences #15710) for 10 minutes. Fixed cells were permeabilized with 160 µL of 5% Triton X-100 (MilliporeSigma #T8787-50ML) + RNase Inhibitor for 3 minutes. Permeabilized cells were then centrifuged at 500 g for 5 minutes at 4°C and resuspended in 500 µL of PBS + RI and mixed with an additional 500 µL of 100 mM Tris-HCL (ThermoFisher #AM9855G) and then 20 µL 5% Triton X-100. Cells were then centrifuged at 500 g for 5 minutes at 4°C and the pellet resuspended in 300 µL 0.5X PBS + RI, then passed through a 40 µm filter. Filtered cells were counted using a hemocytometer and diluted to 1×10^6^ cells/mL in 0.5X PBS + RI. Labeled tubes were placed in a RT (room temperature) Mr. Frosty (Thermo Fisher #5100-0001) and placed into a -80°C freezer for storage.

In-cell reverse transcription, ligation barcoding, lysis, and library preparation were carried out according to the protocol described in Rosenberg and Roco et al.^35^. Vials were thawed by placing tubes in a 37°C water bath, and vial contents were pipetted into wells containing barcoded, well-specific reverse transcription primers and a reverse transcription reaction mix. Each well contained a mixture of random hexamer and anchored poly(dT)_15 barcoded RT primers. Two different sequencing experiments were conducted. Sequencing Experiment 1 contained samples across Day 12/14 and Day 24/26, while Sequencing Experiment 2 contained samples across D0-Stem Cells, Day 90, and one re-sequenced sample each of Day 12 and Day 24 that had been included in Sequencing Experiment 1 to control for sequencing batch variability. Sample metadata including sequencing experiments can be found in **Supplementary Table S1**. For Sequencing Batch 1 (48 samples; see **Supplementary Table S1**), each sample was placed in a single well (4,000 cells per input sample) for a total of 192,000 cells. For Sequencing Batch 2 (9 samples, see **Supplementary Table S1**), each D0 and D90 sample was split over 6 wells (24,000 input cells per sample). The D12 and D24 samples in Sequencing Batch 2 were each split into 3 wells (12,000 input cells per sample). In both sequencing batches, after three rounds of barcoding, the cells were counted and divided into sub-libraries of 5000 cells before lysis. These sub-libraries were barcoded with a fourth unique barcode each and processed for sequencing on an Illumina NextSeq.

### scRNA-seq data processing

Sequence alignment and quantification of intronic and exonic UMI (unique molecular identifier) counts were performed as described in Rosenberg et al.^35^ using STAR^72^ for alignment, TagReadWithGeneExon from Drop-seq tools to assign reads to genes, and Starcode^73^ to collapse UMIs. Per gene intronic and exonic UMI counts were collapsed prior to generating the count matrix. To filter out ambient RNA barcodes, the number of UMIs per barcode were plotted with barcodes ordered by number of UMIs in descending order. The inflection point (or knee) in the plot was used as the threshold to separate barcodes originating from intact cells and barcodes originating from ambient RNA; all barcodes that fell below this threshold were removed. The remaining cells were further filtered for mitochondrial content (percent of UMI counts coming from mitochondrial genes) to remove low quality cells and total UMI counts to remove potential doublets with very large counts. The two sequencing batches were filtered separately. For Sequencing Batch 1 (D12, D14, D24, D26; see **Supplementary Table S1**), cells within the highest 5% of percent mitochondrial and total UMI counts distributions were removed. For Sequencing Batch 2 (D0, D90; see **Supplementary Table S1**), D90 cells had higher percentage mitochondrial UMIs than D0, which is consistent with previous studies showing an increase in mitochondrial content and number during cardiomyocyte maturation^74,75^. D90 cells also had a decrease in total UMI counts compared to D0, which is consistent with other studies indicating a decrease after differentiation^76^. Prompted by these observations, we used the same filtering approach (removing the highest 5% of cells based on UMI and mitochondrial distributions) but applied it independently to each time point (D0, D12, D24 and D90), which led to different maximum cutoffs being used for each time point. For both sequencing batches, genes detected in less than 10 cells and cells with less than 200 transcribed genes were removed from the matrix prior to clustering and visualization.

### scRNA-seq clustering and visualization of D0 and Protocol 1 D12, D24, and D90 samples

Cells were clustered and visualized using the R (version 3.5.1) package Seurat (version 2.3.4)^77^. Normalized transcript abundance for each gene was calculated by dividing counts by the total counts per cell, multiplying by a scaling factor (10000) and log transforming the result using log1p (NormalizeData function in Seurat). For stem cell (D0) and Protocol 1 samples (D12, D24, D90), highly variable genes to be used for dimensionality reduction and clustering were identified using FindVariableGenes with the following parameters: mean.function = ExpMean, dispersion.function = LogVMR, x.low.cutoff = 0.05, x.high.cutoff = 4, y.cutoff = 0.5 (these parameters were chosen to define outlier genes after manual inspection of mean expression vs. dispersion plot). Counts were scaled using ScaleData with default parameters without regressing out any variables. Principal component analysis (PCA) was performed on the normalized and scaled matrix (with highly variable genes only) using RunPCA. Standard deviations of the principal components were plotted to determine the number of components to retain for clustering and visualization, and principal components above the inflection point in the plot were retained. Cells were clustered using the Jaccard-Louvain method^78^, which is based on shared nearest neighbor modularity optimization (FindClusters function with resolution=c(0.3, 0.4, 0.5, 0.6, 0.8, 1) and the following standard parameters: algorithm=1 (original Louvain algorithm), modularity.fxn=1 (standard modularity function)). Uniform Manifold Approximation and Projection (UMAP)^79^ was used to visualize cells in a two-dimensional space (RunUMAP function with default parameters). In **Figs. 1-2** and **Supplementary Fig. S2**, where only Protocol 1 is shown (D0, D12, D24, D90; n = 11,619 cells), clustering with resolution 0.5 was used for visualization, differential expression, and other downstream analyses. Pearson correlation was calculated for cardiac troponin T transcript and protein abundance between flow cytometry-based abundance (% cTnT positive cells, described in “Flow cytometry of scRNA-seq samples”) and scRNA-seq based abundance (% of *TNNT2* positive cells; **Fig. 2B**). Flow cytometry and scRNA-seq were performed on different cells obtained from the same differentiation sample (same differentiation well). Plots were created using R package ggplot2 (version 3.3.0)^80^, and violin plots were created using R package scrattch.vis (version 0.0.210)^81^.

### scRNA-seq clustering and visualization of D12, D14, D24, D26 samples

To compare differentiation protocols, cell lines, and differentiation experiments, Protocols 1 and 2 early and intermediate time point cells (D12, D14, D24, D26; n = 15,878 cells) were clustered independently using the Seurat workflow described above (clustering with resolution 0.4 is shown, and clusters are numbered c0-c10; **Fig. 4**). Pairwise Spearman rank correlation coefficients were calculated between the three cell lines, five differentiation experiments, and two differentiation protocols (**Fig. 5A, Supplementary Fig. S6B, S7A**). For independent clustering of time points and differentiation experiments, clustering with resolution 0.4 is shown in **Fig. 5C-G, Supplementary Fig. S6C-E, Supplementary Fig. S7C-G**). Per gene variance explained (R^2^/coefficient of determination) for variables of interest (day of differentiation, protocol, differentiation experiment, differentiation protocol, cell line, # of genes detected, # of UMIs detected) was calculated using getVarianceExplained in R package scater (version 1.14.6)^82^. Distribution of variance explained for variables of interest is plotted for the top 5% of highly variable genes (602 genes identified using getTopHVGs from R packager scran version 1.14.6)^83^ in the D12, 24, 24, and 26 population.

### scRNA-seq differential expression analysis

Differentially expressed (DE) genes between clusters were identified by performing pairwise comparisons with edgeR (version 3.26.0)^84,85^. RNA composition normalization was performed with calcNormFactors, and negative binomial dispersions were estimated with estimateDisp. DE genes were identified by fitting gene-wise generalized linear models (glmFit; did not include intercept term in design) and performing a likelihood ratio test (glmLRT with contrasts). We retained genes with an absolute log2 fold change >= 1 and Benjamini-Hochberg adjusted p-value < 0.05). To remove potential false positives caused by dropouts in low depth data (i.e., gene expressed very highly in only a few cells within a cluster), we calculated the fraction of cells positive for each gene and retained only genes where the up-regulated group had at least 30% of cells positive for that gene. Heatmaps of top differentially expressed genes (**Fig. 2C, Fig. 5E, Supplementary Fig. S2A, Supplementary Fig. S7G**) were created using R package pheatmap (version 1.0.12)^86^ and show scaled transcript abundance (normalized counts for each gene were scaled and centered). Dendrograms in heatmaps show hierarchical clustering of genes. For visualization purposes, maximum scaled transcript abundance cutoffs were applied in some cases (see heatmap figure legends), and cells are grouped by cluster. Biological Process (BP) gene ontology (GO) enrichment analysis was performed on differentially expressed genes between the following clusters: 1) C2/D0 + C0/D12, 2) C0/D12 + C1/D24, and 3) C1/D24 + C3/D90 using enrichGO function in the R package clusterProfiler (version 3.14.0)^87^. Redundancy in enriched GO terms was removed using the simplify function in clusterProfiler, and the top 10 simplified terms from each pairwise comparison are visualized in Supplementary Fig. S3C. All statistical calculations were performed in R 3.5.1, and plotting was performed using ggplot2 (version 3.3.0)^80^.

### scRNA-seq feature selection analysis

In addition to differential expression analysis based on pairwise cluster comparisons, an orthogonal clustering-independent method was used for identifying genes that change between early and intermediate differentiation time points (D12 and D24) and intermediate and late differentiation time points (D24 and D90). We fit generalized linear models with an elastic net penalty using the glmnet R package (version 2.0-13)^30,31^ with alpha = 0.5 and a sequence of 100 values of lambda, the regularization parameter. The response variable was time point (D12 vs. D24 and D24 vs. D90 for Protocol 1 and D14 vs. D26 for Protocol 2). The input count matrix was filtered down to gene symbols that had a corresponding Entrez Gene ID, and cells were filtered to include only cardiomyocytes (cells positive for cardiomyocyte marker gene *TNNT2*). Cells were split into a training set (90% of cells; for Protocol 1 D12 vs. D24, n = 4,283 D12 cells and n = 2,014 D24 cells; for Protocol 1 D24 vs. D90, n = 2,016 D24 cells and n = 2,262 D90 cells; for Protocol 2 D14 vs. D26, n = 4,228 D14 cells and n = 2,290 D26 cells) and a test set (10% of cells; for Protocol 1 D12 vs. D24, n = 481 D12 cells and n = 227 D24 cells; for Protocol 1 D24 vs. D90, n = 227 D24 cells and n = 253 D90 cells; for Protocol 2 D14 vs. D26, n = 476 D14 cells and n = 259 D26 cells). 1000 bootstrap rounds (sampled 80% of cells without replacement at each round) were run with the training set to identify genes that had non-zero coefficients in all 1000 rounds at different values of the regularization parameter, lambda. Genes with non-zero coefficients in all 1000 bootstrap rounds at a given value of the regularization parameter, lambda, were reported as selected (see **Supplementary Table S3** for top 40 selected genes ranked by lambda when first selected. Each set of genes selected from Protocol 1 (D12 vs. D24, D24 vs. D90) was used to predict the time point for cells in the corresponding test set by fitting a binomial generalized linear model (glmnet with ridge penalty, alpha = 0, and lambda = 1×10^−6^) with the training data set and then using the model to predict time point in the test data set (**Supplementary Fig. S3**). Accuracy of prediction was calculated using the confusionMatrix function from the R package caret (version 6.0-85)^88^ with positive = D12 time point for D12 vs D24, and positive = D24 time point for D24 vs D90. As a control for the accuracy of each selected gene set size, we took 100 random gene samples of the same size and calculated accuracy of predicting time point in the test data set. Accuracies are shown as box plots (outliers hidden) for random gene sets and as individual points for selected gene sets (**Fig. 3B** for D12 vs. D24, **Supplementary Fig. S4D** for D24 vs. D90).

### Replating cardiomyocytes for imaging and RNA FISH

After performing cardiomyocyte differentiation, cells were dissociated into single cells at D12 (described in **“**Single cell dissociation for scRNA-seq” section) and seeded into glass bottom multiwell plates with an aliquot being used for flow cytometry analysis as described above. First, glass bottom multiwell plates (24-well, Cellvis P24-1.5H-N) were incubated at RT with 0.5 M glacial acetic acid (Fisher Scientific #BP1185-500) for 20–60 min and washed once with sterile milliQ (MQ) water. Wells were then treated with 0.1% PEI (Sigma Aldrich #408727-100ML) solution in sterile MQ water for 16–72 h at 4°C and rinsed with DPBS and sterile MQ water. Wells were then incubated with 25 µg/mL natural mouse laminin (Gibco #23017-015) diluted in sterile MQ water overnight at 4°C and removed immediately preceding cell plating. Cells were seeded at a density of 35,000 to 50,000 cells per well in RPMI-1640 supplemented with B27 containing insulin, 1% P/S, and 10 µM Rock Inhibitor. Media was changed after 24 hrs to RPMI-1640 supplemented with B27 containing insulin and 1% P/S, and media was changed every 2-3 days after until fixation. Cells were fixed at time points indicated in text for RNA FISH; the D18 time point reflects single cell dissociation at D12 and a 6-day recovery period after replating, and the D30 time points reflect additional maturation. Cells at the D30 time point were fixed between D29-D30. Both time points (D18 and D30) were seeded from the same source population of D12 differentiated cardiomyocytes that were replated together and maintained in parallel until fixation. Cells were fixed by removing media and washing twice with RNAse-free PBS, then incubated for 10 min at RT in a 4% paraformaldehyde solution (Electron Microscopy Sciences #15710). Fixation solution was removed, wells were washed once more with RNAse-free PBS, then stored in 70% ethanol at -20°C until RNA hybridization was performed.

### RNA FISH using HCR v3.0

Gene validation by RNA FISH was performed using the HCR v3.0 method, following the HCR v3.0 protocol for “Mammalian cells on a slide” with modifications to adapt for samples on glass bottom multiwell plates (Molecular Instruments, https://www.molecularinstruments.com/protocols), as previously described^65^. Fixed wells were washed 4 times with 500 µL 2X SSC (Invitrogen #15557-044), incubated for 30-60 min at 37°C in probe hybridization buffer (Molecular Instruments), and hybridized overnight with 1.2 pmol of each probe set mixture containing 400 U/mL RNAse inhibitor (Enzymatics #Y9240L) at 37°C (250 µL per well). Custom probe sets can be found in **Supplementary Table S5**. Wells were probed for three genes in each experiment. Wells were washed with probe wash buffer (Molecular Instruments) supplemented with 400 U/mL RNAse inhibitor at 37°C for 30 min, washed 4 times with 2X SSC at RT, and incubated in amplification buffer (Molecular Instruments) for 30-60 min at RT. Hairpin amplifiers were prepared during this time; 18 pmol of hairpin amplifiers were heated to 95°C for 90 sec, protected from light and cooled, then combined and added into an amplification buffer containing 400 U/mL RNAse inhibitor. Hairpin mixtures were added to the appropriate wells (250 µL per well) and incubated for 45 min at RT, protected from light. Hairpin solution was removed, wells were washed with 2X SSC four times, nuclei were labeled with 2 µg/mL DAPI in 2X SSC for 5 min, followed by additional washes with 2X SSC. Samples were stored protected from light at 4°C in 2X SSC with 400 U/mL RNAse inhibitor until imaging.

### Imaging RNA FISH samples

Imaging cardiomyocytes after RNA FISH (**Fig. 1E, 3E, Supplementary Fig. 5**) was performed on a Zeiss spinning-disk microscope with a 40x/1.2 NA W C-Apochromat Korr UV-vis infrared (IR) objective (Zeiss) and a 1.2x tube lens adapter for a final magnification of 48x, a CSU-X1 spinning-disk head (Yokogawa), and Orca Flash 4.0 camera (Hamamatsu) (pixel size 0.271µm in X-Y after 2×2 binning and 0.29 µm in Z). Standard laser lines (405, 488, 561, 640 nm), primary dichroic (RQFT 405, 488, 568, 647 nm) and the following Band Pass (BP) filter sets (Chroma) were used for fluorescent imaging: 450/50 nm for detection of DAPI, and 525/50 nm, 600/50 nm, and 690/50 nm for detection of RNA FISH probes. Brightfield images were acquired using an LED light source with peak emission of 740 nm with narrow range and a BP filter 706/95 nm for brightfield light collection.

### Manual cell annotations in Napari

Multi-channel Z-stacks were loaded into Napari (https://napari.org)^89^ and 2D single cell masks were generated by manually drawing cell boundaries in 2D while incorporating information from all channels collected during imaging (brightfield, two FISH probe channels, nuclei via DNA dye (DAPI), and alpha-actinin-2-mEGFP (structure) signal if present). Single cell masks in fields of view (FOVs) were hand drawn by a single human expert. Cell boundaries were manually drawn for cells that were mostly within the FOV, and low-confidence/high cell density regions with many overlapping cells were avoided. 2D single cell masks were used downstream to create single cell transcript abundance measurements.

### DNA (Nuclear) segmentation

Nuclear segmentation in 2D using the DNA channel was performed using CellProfiler (version 3.1.8)^90^. See CellProfiler pipeline for Zeiss in https://github.com/AllenCellModeling/fish_morphology_code for details. The DNA channel maximum intensity projection (MIP) was normalized with CellProfiler’s RescaleIntensity module from the 5^th^ percentile to 95^th^ percentile of the raw image. Nuclei were segmented using minimum cross entropy thresholding to define the probability distributions of foreground and background regions in an image using CellProfiler’s IdentifyPrimaryObjects module. Clumped objects were filtered by shape to identify nuclear objects in close proximity. Objects smaller than 500 pixels were considered debris and discarded. Nuclei were assigned to a cell if their centroids fell within the 2D segmented cell object. Unassigned nuclear objects were discarded and not used for further analysis.

### RNA spot segmentation and feature extraction

RNA FISH transcripts were segmented using a transcript-specific segmentation workflow in the Allen Cell Structure Segmenter (Allen Cell Explorer)^91^. MIP image intensities were normalized, a Gaussian smoothing filter was applied to all images, and a 2D spot filter algorithm was applied to segment the transcript signal, where each transcript signal represented the location of the RNA as a diffraction-limited spot. This filter accounted for both the dot radius and the filter response to generate a binary result. Dot intensity levels varied by transcript and thus the filter response parameter was optimized for each RNA species, as listed in the table below. Segmented RNA FISH spots were quantified for each manually segmented cell in an FOV using CellProfiler’s IdentifyPrimaryObjects and MeasureObjectSizeShape modules.

| Microscope | Probe set | Filter response parameter |
| --- | --- | --- |
| Zeiss | *TNNT2*-488 | 0.02 |
|  | *TNNT2*-638 | 0.03 |
|  | *MYL2*-488 | 0.06 |
|  | *MYL2*-638 | 0.05/0.06 |
|  | *MYL7*-561 | 0.01 |
|  | *MYH6*-488 | 0.01 |
|  | *MYH6*-561 | 0.02 |
|  | *MYH7*-561 | 0.01 |
|  | *MYH7*-638 | 0.02 |
|  | *ESRRG*-638 | 0.03 |
|  | *MEIS2*-638 | 0.01 |
|  | *FABP3*-638 | 0.04 |
|  | *H19*-561 | 0.01 |
|  | *VCAN*-561 | 0.02 |
|  | *COL2A1*-488 | 0.04 |

All data used to generate figures in this manuscript is available at: https://open.quiltdata.com/b/allencell/packages/aics/wtc11_hipsc_cardiomyocyte_scrnaseq_d0_to_d90

Code used for analysis and to generate figures is available at: https://github.com/AllenCell/cardio_scrnaseq_paper_code

### Statistical Analysis

Details of specific statistical analyses for each section, sample sizes, and statistical tests used are given in the Methods and in the corresponding figure legends.

