## Supplementary Materials for "A comprehensive analysis of gene expression changes in a high replicate and open-source dataset of differentiating hiPSC-derived cardiomyocytes"

Supplementary Figure S1

**A**

| | D0 (undifferentiated) | | | Protocol 1 D12 | | | Protocol 1 D24 | | | Protocol 1 D90 | | | Protocol 2 D14 | | | Protocol 2 D26 | | |
|---|---|---|---|---|---|---|---|---|---|---|---|---|---|---|---|---|---|---|
| | WTC-11 | *TOMM20* | *TNNI1* | WTC-11 | *TOMM20* | *TNNI1* | WTC-11 | *TOMM20* | *TNNI1* | WTC-11 | *TOMM20* | *TNNI1* | WTC-11 | *TOMM20* | *TNNI1* | WTC-11 | *TOMM20* | *TNNI1* |
| Diff Exp1 | | | | 1 | | 1 | | | | | | | 1 | | 1 | | | |
| Diff Exp2 | | | | 1 | 1 | 1 | | | | | | | 1 | 1 | 1 | | | |
| Diff Exp3 | | | | | 1 | 1 | 1 | | 1 | | | | | 1 | 1 | 1 | | 1 |
| Diff Exp4 | | | | | 2 | 1 | | 2 | 1 | | | | | 2 | 1 | | 2 | 1 |
| Diff Exp5 | | | | 1 | 2 | 1 | 1 | 1 | 2 | | | | 1 | 2 | 1 | 3 | 1 | 2 |
| Diff Exp6 | | | | | | | | | | 2 | | 1 | | | | | | |
| Diff Exp7 | | | | | | | | | | | | 2 | | | | | | |
| Undiff | 1 | | 1 | | | | | | | | | | | | | | | |

**B**

| | D0 (undifferentiated) | | | Protocol 1 D12 | | | Protocol 1 D24 | | | Protocol 1 D90 | | | Protocol 2 D14 | | | Protocol 2 D26 | | |
|---|---|---|---|---|---|---|---|---|---|---|---|---|---|---|---|---|---|---|
| | WTC-11 | *TOMM20* | *TNNI1* | WTC-11 | *TOMM20* | *TNNI1* | WTC-11 | *TOMM20* | *TNNI1* | WTC-11 | *TOMM20* | *TNNI1* | WTC-11 | *TOMM20* | *TNNI1* | WTC-11 | *TOMM20* | *TNNI1* |
| Diff Exp1 | | | | 1 | | 1 | | | | | | | 1 | | 1 | | | |
| Diff Exp2 | | | | 1 | 1 | 1 | | | | | | | 1 | 1 | 1 | | | |
| Diff Exp3 | | | | | 1 | 1 | 1 | | 1 | | | | | 1 | 1 | 1 | | 1 |
| Diff Exp4 | | | | | 2 | 1 | | 2 | 1 | | | | | 2 | 1 | | 2 | 1 |
| Diff Exp5 | | | | 1 | 2 | 1 | 1 | 1 | 2 | | | | 1 | 2 | 1 | 3 | 1 | 2 |
| Diff Exp6 | | | | | | | | | | 2 | | 1 | | | | | | |
| Diff Exp7 | | | | | | | | | | | | 2 | | | | | | |
| Undiff | 1 | | 1 | | | | | | | | | | | | | | | |

**C**

| | D0 (undifferentiated) | | | Protocol 1 D12 | | | Protocol 1 D24 | | | Protocol 1 D90 | | | Protocol 2 D14 | | | Protocol 2 D26 | | |
|---|---|---|---|---|---|---|---|---|---|---|---|---|---|---|---|---|---|---|
| | WTC-11 | *TOMM20* | *TNNI1* | WTC-11 | *TOMM20* | *TNNI1* | WTC-11 | *TOMM20* | *TNNI1* | WTC-11 | *TOMM20* | *TNNI1* | WTC-11 | *TOMM20* | *TNNI1* | WTC-11 | *TOMM20* | *TNNI1* |
| Diff Exp1 | | | | 1 | | 1 | | | | | | | 1 | | 1 | | | |
| Diff Exp2 | | | | 1 | 1 | 1 | | | | | | | 1 | 1 | 1 | | | |
| Diff Exp3 | | | | | 1 | 1 | 1 | | 1 | | | | | 1 | 1 | 1 | | 1 |
| Diff Exp4 | | | | | 2 | 1 | | 2 | 1 | | | | | 2 | 1 | | 2 | 1 |
| Diff Exp5 | | | | 1 | 2 | 1 | 1 | 1 | 2 | | | | 1 | 2 | 1 | 3 | 1 | 2 |
| Diff Exp6 | | | | | | | | | | 2 | | 1 | | | | | | |
| Diff Exp7 | | | | | | | | | | | | 2 | | | | | | |
| Undiff | 1 | | 1 | | | | | | | | | | | | | | | |

**D**

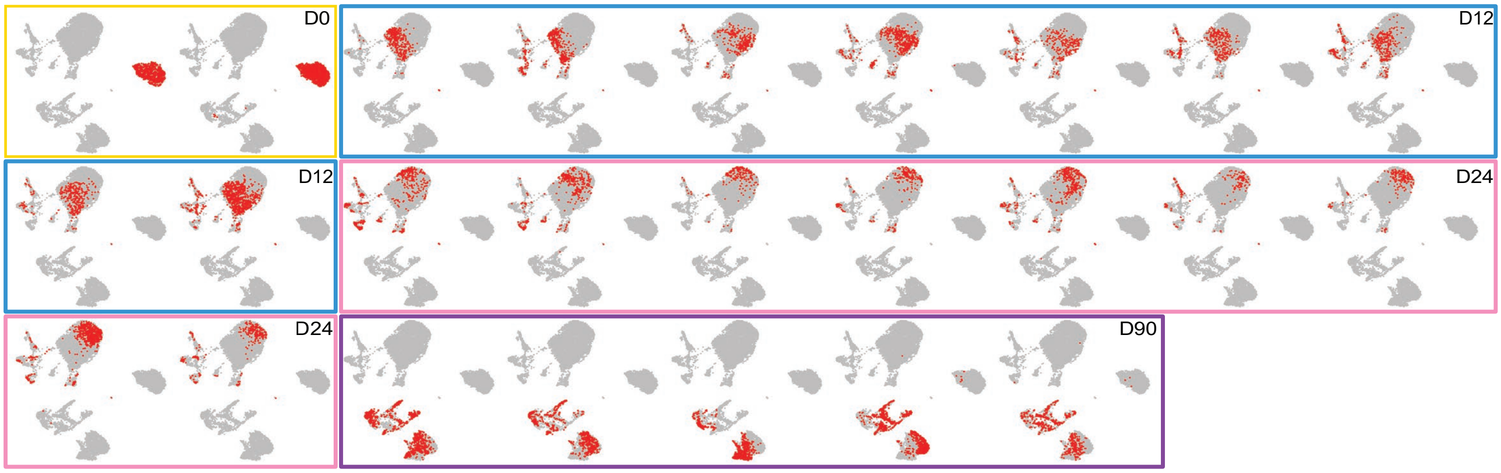


**E**

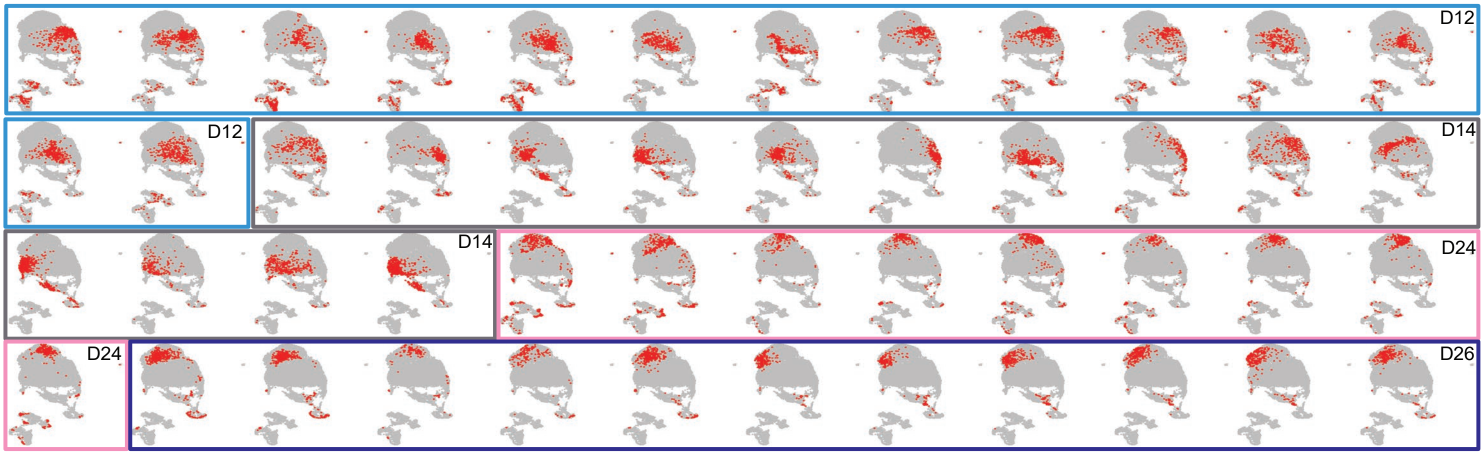

**Supplementary Figure S1. Sample overview**

A. Table summarizing all scRNA-seq samples. Each row represents one differentiation experiment (plates/samples set up on one day; see Fig. 5B). Numbers in the table indicate the number of wells that were collected. Wells were not pooled.

B. Table highlighting scRNA-seq samples used in Figs. 1-3, Supplementary Figs. S2-S4, focusing on gene expression during cardiomyocyte differentiation with Protocol 1 at D0, D12, D24 and D90.

C. Table highlighting scRNA-seq samples used in reproducibility analysis (Figs. 4-5, Supplementary Figs. S6-S7).

D. UMAPs showing each of the samples from D0, D12, D24, D90 Protocol 1 analysis (D0 n = 2 samples; D12 n = 9 samples; D24 n = 9 samples; D90 n = 5 samples). Each individual sample highlighted in green in B is shown in a separate UMAP in red. The UMAP is the same as in Fig. 1. Box colored by time point (colors as in Fig. 1D) is drawn around all samples from a given time point.

E. UMAPs showing each of the samples from D12, D14, D24, D26 reproducibility analysis (D12 n = 14 samples; D14 n = 14 samples; D24 n = 9 samples; D26 n = 11 samples). Each individual sample highlighted in green in C is shown in a separate UMAP in red. The UMAP is the same as in Fig. 4. Box colored by time point (colors as in Fig. 4C) is drawn around all samples from a given time point.

Supplementary Figure S2

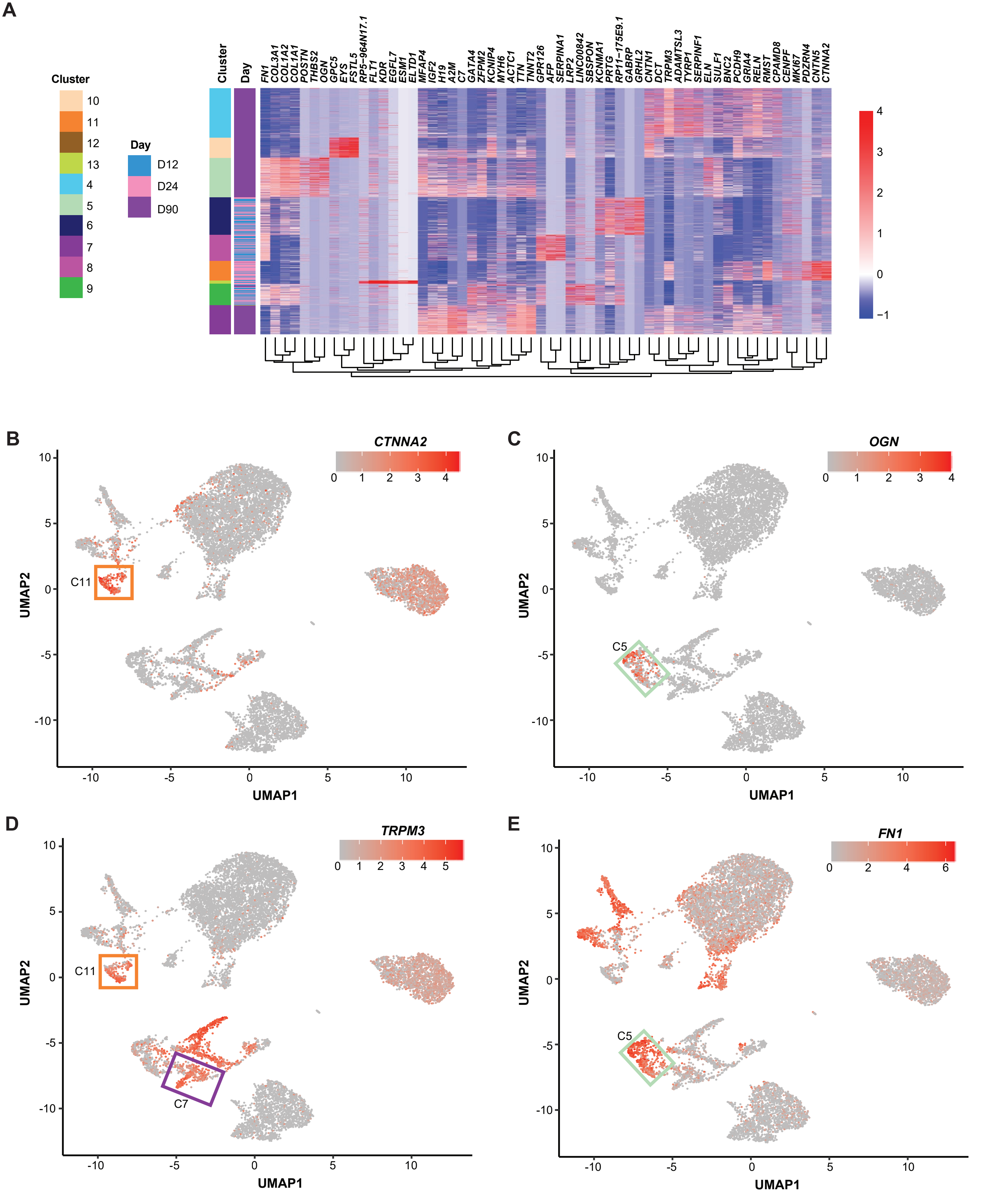

**Supplementary Figure S2: Non-cardiomyocyte cell types identified in D12, D24, and D90 samples generated with differentiation Protocol 1**

A. Heatmap showing top differentially expressed genes from each pairwise cluster comparison between pairs of non-cardiomyocyte clusters (non-cardiomyocytes were identified by the absence of marker *TNNT2;* see Fig. 1C-D). Heatmap includes cardiomyocyte cluster 7, proliferative cardiomyocyte cluster 12, and all differentiated non-cardiomyocyte clusters. Normalized transcript abundance was centered and scaled across each gene (z-score color scale to the right of heatmap; red = standard deviations above mean; blue = standard deviations below mean; white = mean; for visualization purposes, 4 was set as the maximum z-score, and z-scores > 4 were set to 4). The dendrogram is based on hierarchical clustering of genes. Each row corresponds to one cell.

B. UMAP from Fig. 1 colored by transcript abundance of *CTNNA2*, highlighting non-cardiomyocyte cluster C11 (orange shading in A and outline on UMAP). Increased red shading reflects higher levels of transcript.

C. UMAP from Fig. 1 colored by transcript abundance of *OGN*, highlighting non-cardiomyocyte cluster C5 (mint green shading in A and outline on UMAP).

D. UMAP from Fig. 1 colored by transcript abundance of *TRPM3*, highlighting non-cardiomyocyte cluster C11 (orange shading in A and outline on UMAP) and cardiomyocyte cluster C7 (purple shading in A and outline on UMAP).

E. UMAP from Fig. 1 colored by transcript abundance of *FN1*, highlighting non-cardiomyocyte cluster C5 (mint green shading in A and outline on UMAP).

Supplementary Figure S3

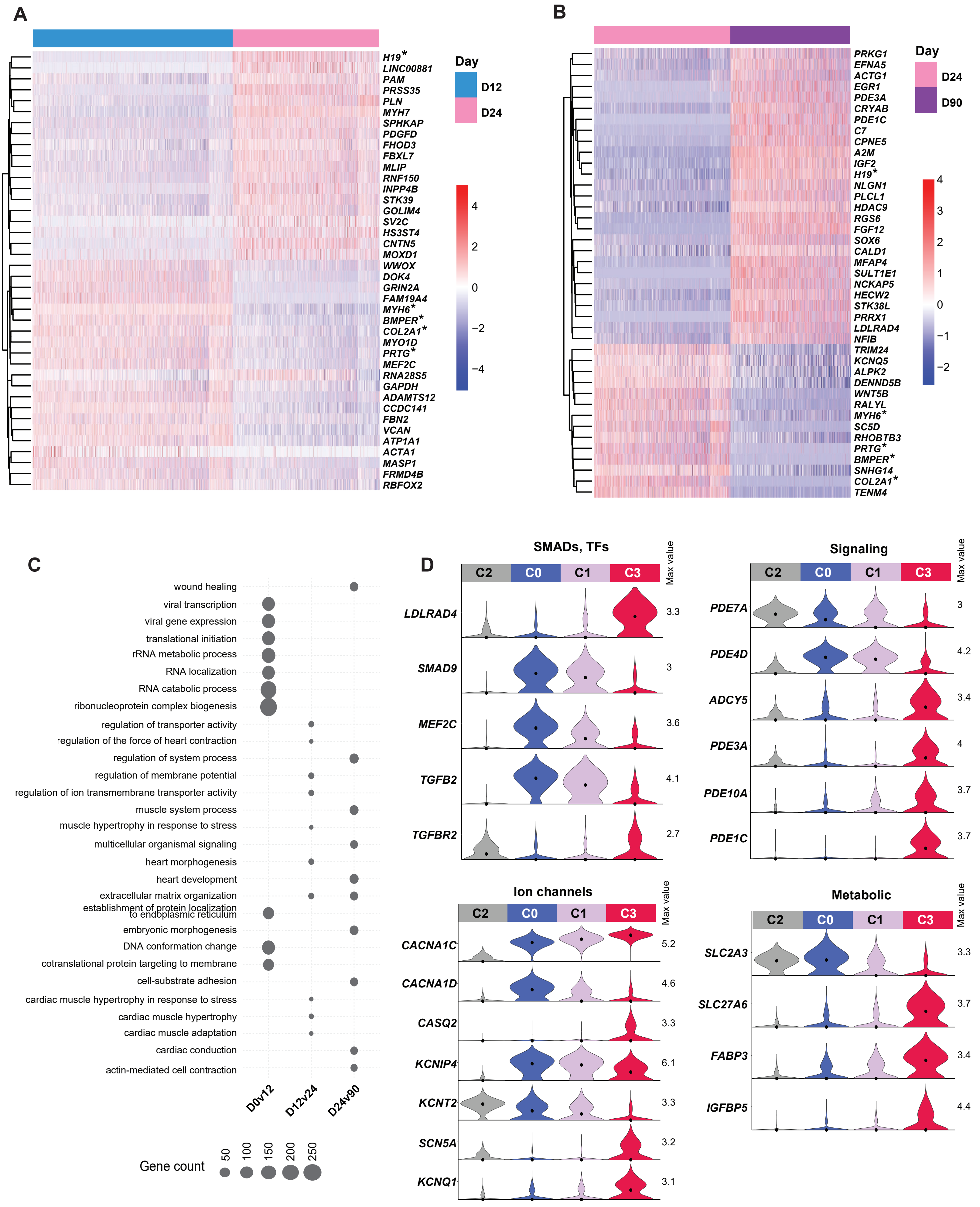

**Supplementary Figure S3: Gene changes in cardiomyocyte populations over time**

A. Heatmap of the top 40 ranked genes from feature selection analysis between D12 and D24 cardiomyocytes. Genes that overlap between (A) and (B) are marked with an "*".
B. Heatmap of the top 40 ranked genes from feature selection analysis between D24 and D90 cardiomyocytes. Genes that overlap between (A) and (B) are marked with an "*".
C. Enriched gene ontology (GO) categories were identified for differentially expressed genes between Day 0 (C2) and Day 12 (C0), Day 12 (C0) and Day 24 (C1), and Day 24 (C1 and Day 90 (C3). Size of the circle represents the number of differentially expressed genes in each category. Top 10 enriched GO categories (ranked by multiple testing adjusted p-value) are shown for each pairwise comparison. See Supplemental Table S4.
D. Functional gene categories that change between D24 and D90 cardiomyocytes. Transcript abundance distributions are shown for C2 (D0), C0 (D12), C1 (D24), and C3 (D90). Max value = maximum value of log1p normalized counts; dot = median.

Supplementary Figure S4

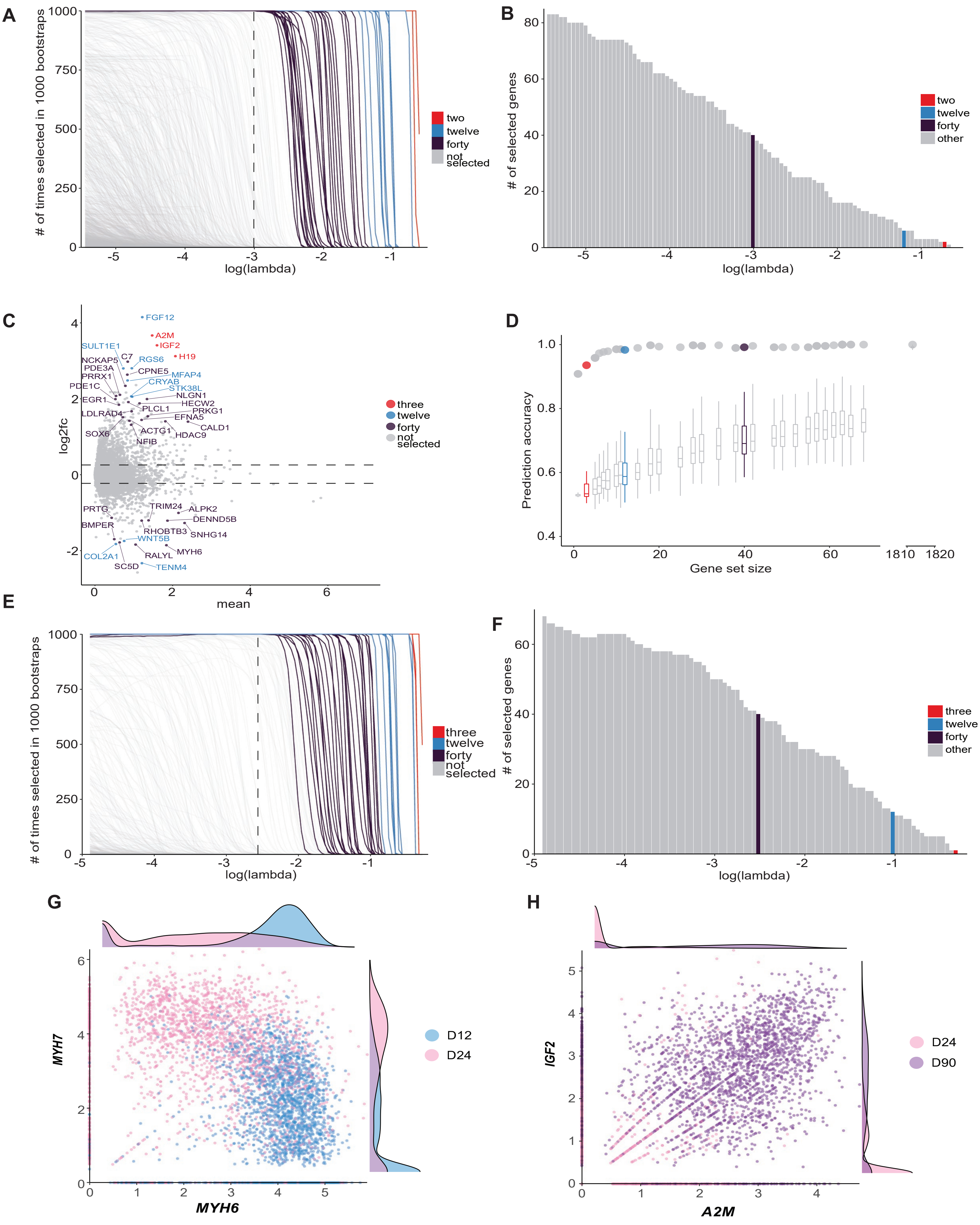

**Supplementary Figure S4: Feature selection analysis in D12 vs D24 and in D24 vs D90 cardiomyocytes**

A. D12 vs. D24 bootstrapped penalized regression selection frequency for each gene across regularization parameter, lambda, sequence. Selected genes are color coded based on size of the selected gene set as in Fig. 3A-B.

B. Number of selected genes from D12 vs. D24 feature selection at different values of lambda. See Fig. 3A-B.

C. D24 vs. D90 cardiomyocyte log2 fold change vs. mean expression. Top feature selected genes are color coded in red (top 3 genes), blue (top 12 genes), and purple (top 40 genes); all other genes are shown in gray.

D. Accuracy of predicting cell age (D24 vs. D90) in holdout data using feature selected gene sets of different sizes. The prediction accuracy for a set of highly variable genes between D24 and D90 is shown as the dot to the right of the x-axis break. Prediction accuracies for random gene sets of the same size are shown as box plots with outliers omitted (100 random samples for each gene set size).

E. D24 vs. D90 bootstrapped penalized regression selection frequency for each gene across regularization parameter, lambda, sequence. Selected genes are color coded based on size of the selected gene set as in C.

F. Number of selected genes from D24 vs. D90 feature selection at different values of lambda.

G. Scatter plot of transcript abundances of the top two D12 vs. D24 feature selection genes, *MYH6* and *MYH7*, in D12 and D24 cardiomyocytes.

H. Scatter plot of transcript abundances of the top two D24 vs. D90 feature selection genes, *IGF2* and *A2M*, in D24 and D90 cardiomyocytes.

Supplementary Figure S5

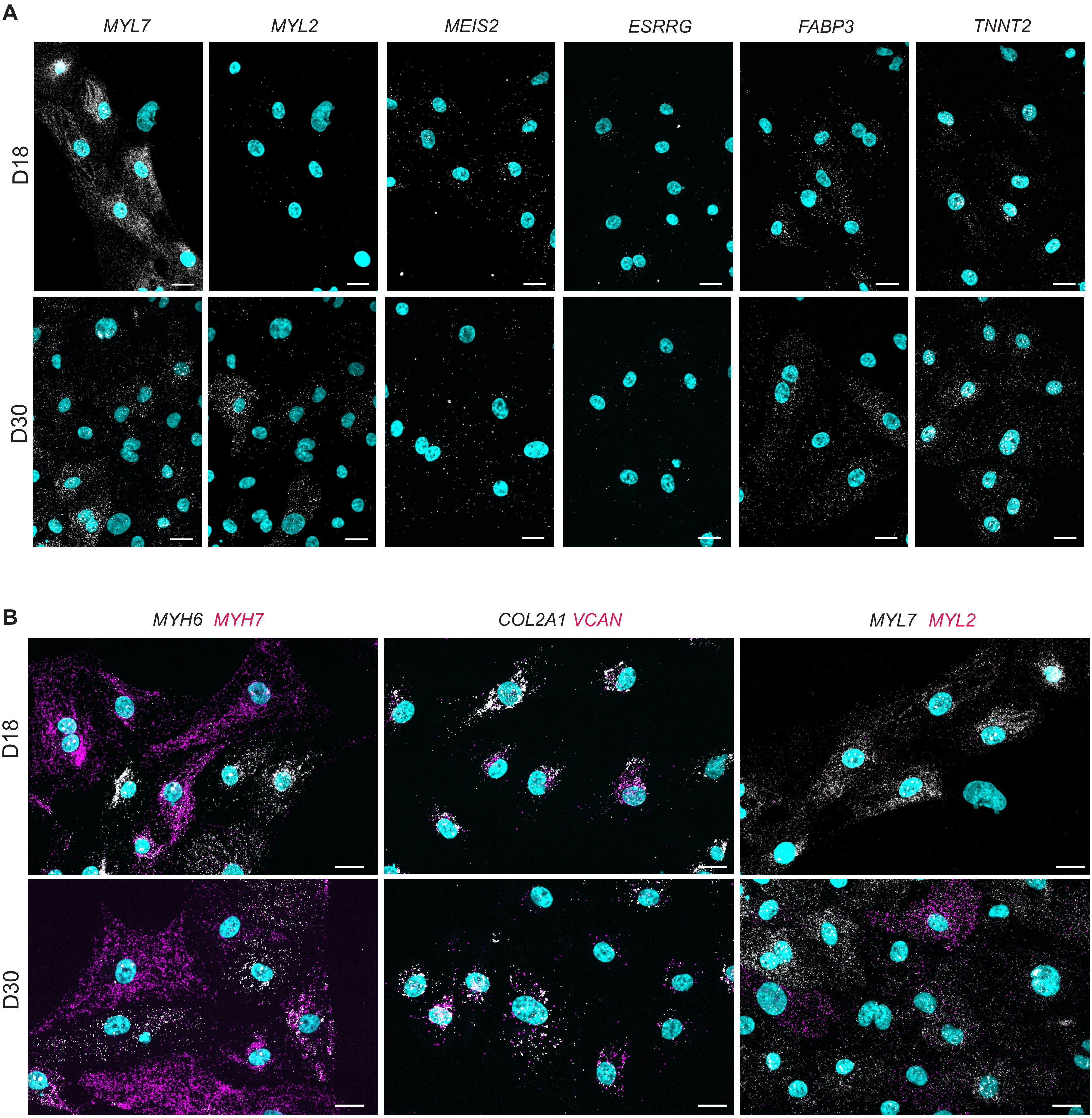

**Supplementary Figure S5: RNA FISH in cardiomyocytes validates genes identified in scRNA-seq data analysis**

A. RNA FISH transcripts are shown in white for the following genes at D18 and D30, as labeled: *MYL7*, *MYL2*, *MEIS2*, *ESRRG*, *FABP3*, *TNNT2*. Nuclei are labeled with DAPI (cyan). Scale bars = 20 μm.

B. RNA FISH transcripts are shown in representative fields of view at D18 and D30 for the following gene pairs: *MYH6* (white) and *MYH7* (magenta), *COL2A1* (white) and *VCAN* (magenta), and *MYL7* (white) and *MYL2* (magenta). Nuclei are labeled with DAPI (cyan). Scale bars = 20 μm. Transcript abundance for all genes shown in this figure is quantified in Fig. 3D.

Supplementary Figure S6

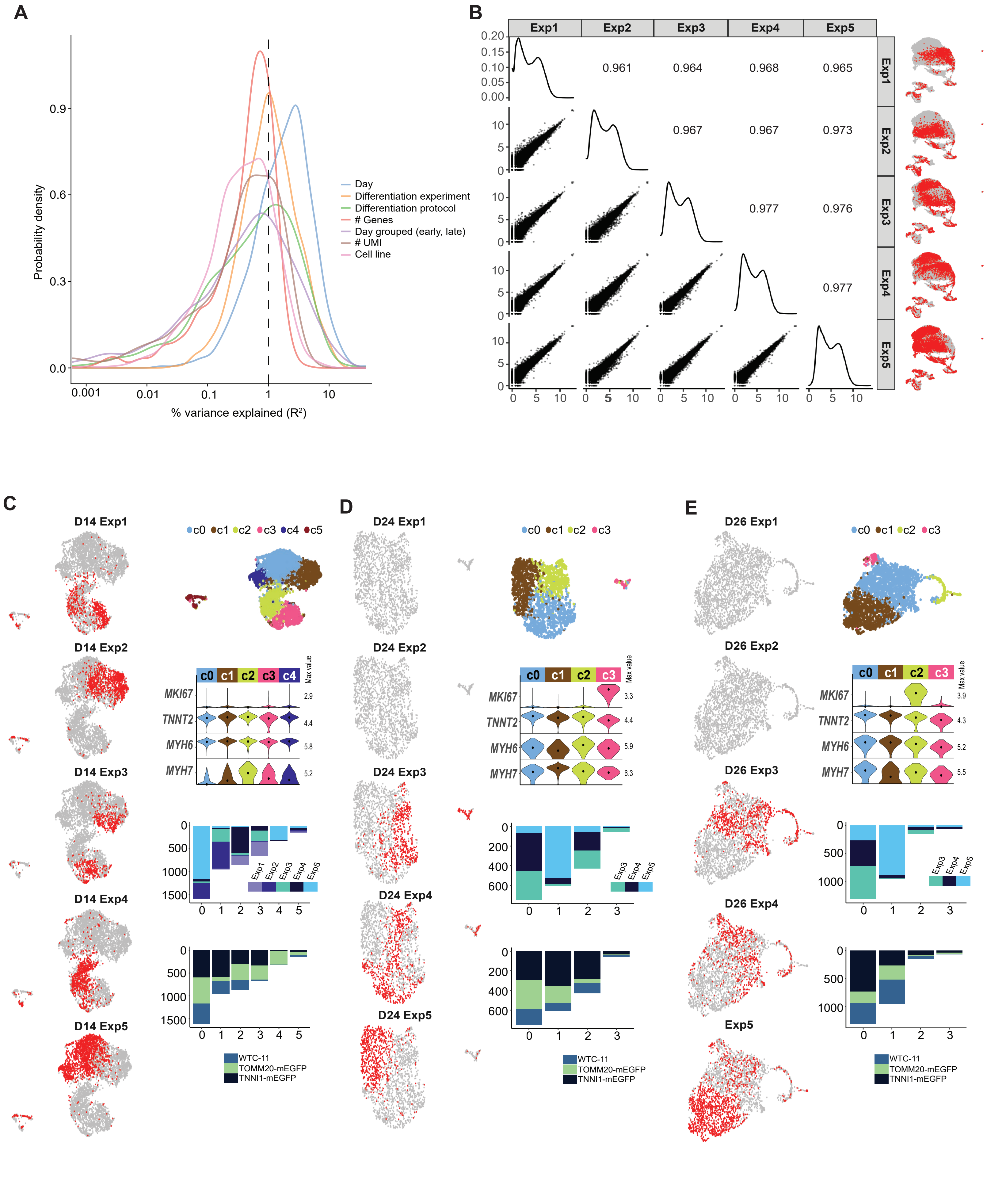

**Supplementary Figure S6: Sources of biological variability and differences in gene expression across differentiation experiments.**

A. Distributions of gene-wise variance explained (coefficient of determination) for cell variables of interest (day of differentiation, differentiation experiment, differentiation protocol, cell line, # of genes detected, # of UMIs detected). Analysis was performed on the set of highly variable genes in D12, D14, D24, and D26 cells.
B. Scatter plots of population transcript abundances between differentiation experiments. Each point is a gene, and Spearman correlations are shown in the upper right. UMAPs on the right highlight cells from each differentiation experiment in red.
C. D14 cardiomyocytes (*TNNT2*+ cells) from all 5 differentiation experiments were independently clustered and visualized using UMAP. Each differentiation experiment is individually colored in red in the left column UMAPs. Top right UMAP is colored by cluster. Violin plot shows distributions of marker genes across clusters with cluster breakdown by differentiation experiment and cell line shown below.
D. D24 cardiomyocytes (*TNNT2*+ cells) from all 5 differentiation experiments were independently clustered and visualized using UMAP. Each differentiation experiment is individually colored in red in the left column of UMAPs. D24 samples were not collected in differentiation experiments 1 and 2. Top right UMAP is colored by cluster. Violin plot shows distributions of marker genes across clusters with cluster breakdown by differentiation experiment and cell line shown below.
E. D26 cardiomyocytes (*TNNT2*+ cells) from all 5 differentiation experiments were independently clustered and visualized using UMAP. Each differentiation experiment is individually colored in red in the left column of UMAPs. D26 samples were not collected in differentiation experiments 1 and 2. Top right UMAP is colored by cluster. Violin plot shows distributions of marker genes across clusters with cluster breakdown by differentiation experiment and cell line shown below.

Supplementary Figure S7

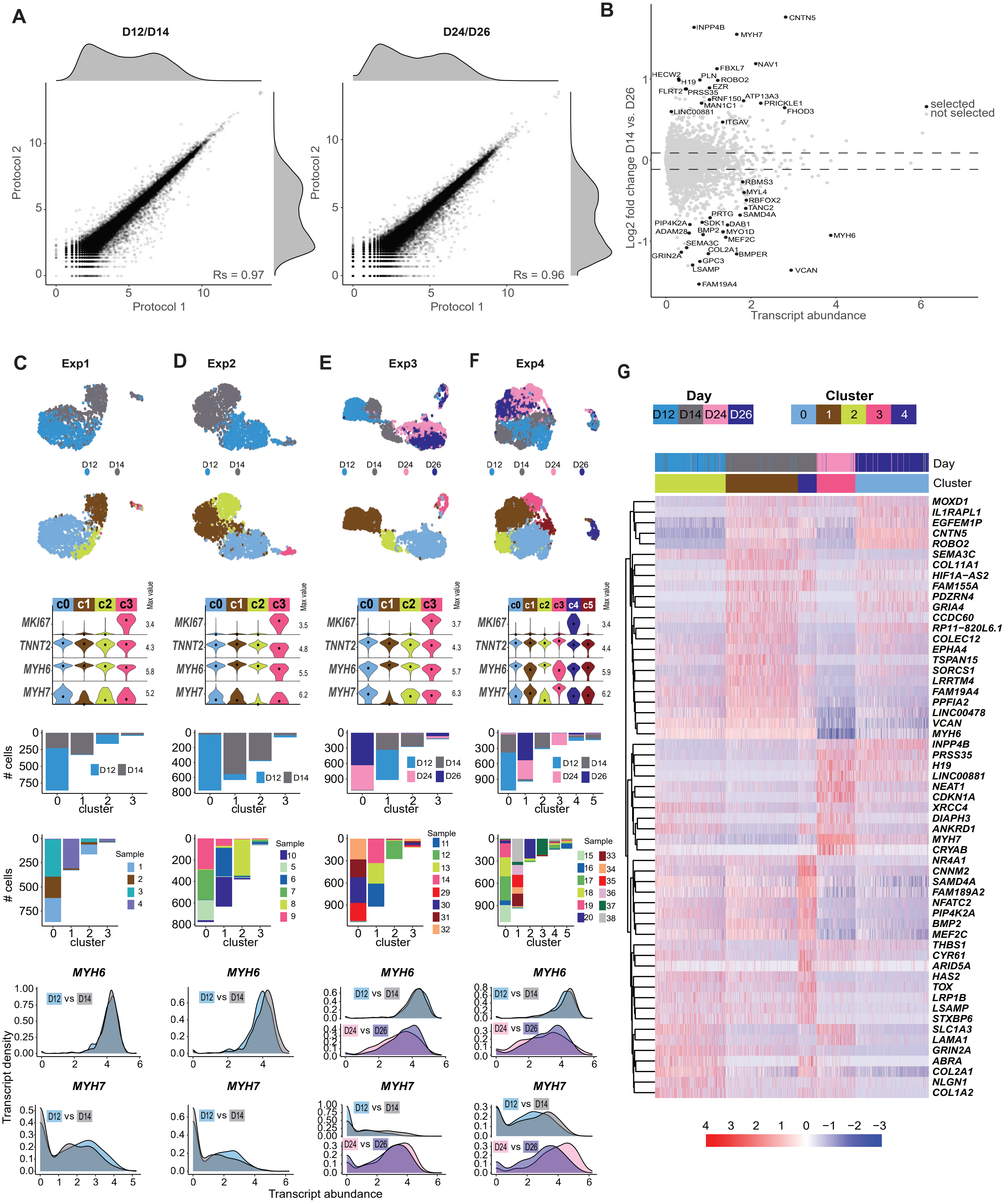

**Supplementary Figure S7: Differences in gene expression between two directed differentiation protocols in D12, 14, 24, and 26 cardiomyocytes**

A. Left: Scatter plot of population transcript abundances between Protocol 1 at D12 and Protocol 2 at D14, respectively. Right: Scatter plot of population transcript abundances between Protocol 1 at D24 and Protocol 2 at D26. Each point represents a gene. Spearman correlation is shown in lower right.

B. Log2 fold change vs. mean expression for Protocol 2 D14 vs D26 cardiomyocytes. Bootstrapped sparse regression analysis was performed as described for Fig. 3A, and selected genes are highlighted in black.

C. Experiment 1 cardiomyocytes (*TNNT2*+ cells) from all collected time points (D12 and D14) were independently clustered and visualized using UMAP to compare differentiation protocols. UMAPs color coded by day/protocol and cluster are shown. Group violin plot shows distributions of marker genes in clusters with cluster breakdown by day and sample shown below. Probability density distribution of *MYH6* and *MYH7* transcript abundance in all Exp1 D12 and D14 cardiomyocytes is shown at the bottom.

D. Experiment 2 cardiomyocytes (*TNNT2*+ cells) from all collected time points (D12 and D14) were independently clustered and visualized using UMAP to compare differentiation protocols. UMAPs color coded by day/protocol and cluster are shown. Group violin plot shows distributions of marker genes in clusters with cluster breakdown by day and sample shown below. Probability density distribution of *MYH6* and *MYH7* transcript abundance in all Exp2 D12 and D14 cardiomyocytes is shown at the bottom.

E. Experiment 3 cardiomyocytes (*TNNT2*+ cells) from all collected time points (D12, D14, D24, and D26) were independently clustered and visualized using UMAP to compare differentiation protocols. UMAPs color coded by day/protocol and cluster are shown. Group violin plot shows distributions of marker genes in clusters with cluster breakdown by day and sample shown below. Probability density distribution of *MYH6* and *MYH7* transcript abundance in all Exp3 D12, D14, D24, and D26 cardiomyocytes is shown at the bottom.

F. Experiment 4 cardiomyocytes (*TNNT2*+ cells) from all collected time points (D12, D14, D24, and D26) were independently clustered and visualized using UMAP to compare differentiation protocols. UMAPs color coded by day/protocol and cluster are shown. Group violin plot shows distributions of marker genes in clusters with cluster breakdown by day and sample shown below. Probability density distribution of *MYH6* and *MYH7* transcript abundance in all Exp4 D12, D14, D24, and D26 cardiomyocytes is shown at the bottom.

G. Heatmap showing top differentially expressed genes between non-proliferative cardiomyocyte clusters (c0-4) in differentiation Experiment 5. See Exp5 clusters in Fig. 5F-G. Normalized transcript abundance was centered and scaled across each row (z-score color scale below heatmap; red = standard deviations above mean; blue = standard deviations below mean; white = mean; for visualization purposes, 4 was set as the maximum z-score, and z-scores > 4 were set to 4). The dendrogram is based on hierarchical clustering of genes. Each column corresponds to one cell.

**Supplementary Table S1: scRNA-seq sample metadata**
Metadata for all samples included in the scRNA-seq data set. Sequencing_batch refers to the sequencing batch that samples belonged to (seq1 or seq2), cell_line indicates one of three cell lines: either AICS0 (AICS-00 WTC-11), AICS11 (AICS-0011 cl.27 TOMM20-mEGFP), or AICS37 (AICS-0037 cl.172 TNNI1-mEGFP). Protocols are listed as Protocol 1 (small molecule), Protocol 1 (small molecule 7.5/7.5), and Protocol 2 (cytokine). See "Cardiomyocyte differentiation using two protocols" section of the Materials and Methods for protocol details. Differentation_experiment refers to independent experiment setups, Exp 1 through Exp7, which correspond to the differentiation_start. Differentiation_start refers to the date at which undifferentiated stem cells were seeded for cardiac differentiation, with the undifferentiated cell passage number and seeding density noted in 10x10^6 cells per well (M, million) in columns H and I, respectively. The harvest date and day when spontaneous beating was observed is recorded (# of days after D0, when differentiation was initiated, see Directed cardiomyocyte differentiation section of Materials and Methods, nr = not recorded). Percent_ctnt is the percent of the harvested population that expressed cardiac troponin T by flow cytometry analysis.

**Supplementary Table S2: Differentially expressed genes between clusters**
List of differentially expressed (DE) genes from pairwise cluster comparisons for D0 (C2), D12 (C0), D24 (C1), and D90 (C3) (see clusters in **Figs. 1, 2, 3, Supplementary Figure S3D**). Each tab is DE genes between one pair of clusters. LogFC = log2 fold change between groups; logCPM = mean log2 of counts per million; LR = likelihood ratio statistics; PValue = p-value before multiple testing correction; FDR = multiple testing adjusted p-values with Benjamini-Hochberg method to control false discovery rate; up = fraction of non-zero cells for gene in up-regulated cluster (cluster in pair with higher transcript abundance of the two); down = fraction of non-zero cells for gene in down-regulated cluster (cluster in pair with lower transcript abundance).

**Supplementary Table S3: Feature selection analysis genes ranked by lambda**
Genes selected from the bootstrapped sparse regression analysis are listed ranked by lambda value (regularization parameter) when they were first selected. Each tab shows a different set (D12 vs D24, D24 vs D90, and D14 vs D26). See **Fig. 3A-B**, and **Supplementary Figs. S4 and S7B**, and scRNA-seq feature selection analysis section of Materials and Methods). Selected = gene with non-zero coefficient in all 1,000 bootstrap rounds at a given value of lambda.

**Supplementary Table S4: Enriched gene ontology categories**
Enriched gene ontology (GO) categories were identified for differentially expressed genes between time points Day 0 (C2) and Day 12 (C0), Day 12 (C0) and Day 24 (C1), and Day 24 (C1) and Day 90 (C3) (group; column J). Table shows top 10 enriched GO categories ranked by adjusted p-value from each pairwise comparison.ID = GO accession; GeneRatio = # of differentially expressed genes that overlap background gene set and are annotated with given GO term / # of differentially expressed genes that overlap background gene set; BgRatio = # of genes from background gene set that are annotated with GO term / # of genes in background gene set; pvalue =hypergeometric p-value; p.adjust = Benjamini-Hochberg adjusted p-value; qvalue = false discovery rate; geneID = list of differentially expressed genes that overlap background gene set and are annotated with given GO term; Count = # of differentially expressed genes that overlap background gene set and are annotated with given GO term. This table is used to make plot in **Supplementary Fig. S3C**.

**Supplementary Table S5: Genes evaluated using RNA FISH**
Genes evaluated using RNA FISH are listed, with protein name (Column B) and NCBI accession number (Column C) for the sequence used to design probe sets listed. All probe sets can be ordered from Molecular Instruments/Molecular Technologies using the unique probe ID listed here (Column D).
